# A new diagnostic modality for bovine tuberculosis: accurate and robust classification of infected cattle using transcriptomics and machine learning

**DOI:** 10.1101/2025.09.01.673540

**Authors:** John F. O’Grady, Adriana Ivich, Gillian P. McHugo, Adnan Khan, Thomas J. Hall, Sarah L. Faherty O’Donnell, Carolina N. Correia, John A. Browne, Valentina Riggio, James G. D. Prendergast, Emily L. Clark, Hubert Pausch, Kieran G. Meade, Isobel C. Gormley, Eamonn Gormley, Stephen V. Gordon, Casey S. Greene, David E. MacHugh

## Abstract

Bovine tuberculosis (bTB) remains recalcitrant to eradication in many endemic countries where current diagnostics are suboptimal. *Mycobacterium bovis* causes bTB and is closely related to *Mycobacterium tuberculosis*, which causes human tuberculosis (hTB). Although blood-based mRNA biomarkers identified through machine learning can discriminate hTB-positive from hTB-negative individuals, similar approaches have not been explored for bTB. Here, we use RNA-seq and machine learning to investigate the utility of blood mRNA as a host-response biomarker for bTB. We identify a 30-gene signature and a 273-gene elastic net classifier that differentiate bTB-positive from bTB-negative cattle, achieving area under the curve (AUC) values of 0.986/0.900 for the former and 0.968/0.938 for the latter in training and testing, respectively. Additionally, we show that these classifiers distinguish bTB-positive cattle from cattle infected with other microbial pathogens (AUC ≥ 0.819). These mRNA-based classifiers represent a promising tool for augmenting current diagnostics to advance global bTB eradication efforts.

## Introduction

*Mycobacterium bovis* causes bovine tuberculosis (bTB), a damaging and endemic disease of cattle that is conservatively estimated to cost the global agricultural industry $3 billion USD annually (*1*). Moreover, *M. bovis* possesses the capacity to infect humans, causing zoonotic tuberculosis (zTB), a disease that is most prevalent in the Global South and due to direct aerosol transmission of the bacillus to animal keepers, or by consumption of contaminated animal by-products (*2–4*). Although variable estimates have been reported for the proportion of human tuberculosis (hTB) cases attributed to *M. bovis* (i.e., zTB), including as high as 28% in Mexico (*5*), the most recent estimates from the World Health Organization (WHO) attributed 140,000 of newly diagnosed active hTB cases (1.4% of 10.0 million) and 11,400 hTB deaths (8.1% of 1.2 million) to zTB (*6*).

In some industrialised countries, such as Ireland, England, and Wales, bTB represents a persistent economic and social problem to the agricultural industry, inflicting combined financial costs in excess of €200 million per year (*7, 8*), and causing significant distress to herdowners (*9*). The high economic cost associated with bTB disease management in these countries stems from state-sponsored bTB eradication programmes. These initiatives, which commenced in the mid-20^th^ century, rely on compulsory skin testing of all herds to identify infected animals that are then removed, slaughtered, and compensated for (*10, 11*). In England and Wales, despite initial success in managing the disease (*12*), the national bTB herd incidence rate in these countries has remained consistently above 6% for the past 15 years (*13*). Similarly, in Ireland, the herd incidence rate fell to a record low of 3.37% in 2015 (*7*); however, since then, the 12-month rolling bTB herd incidence rate has almost doubled, rising to 6.40% by June 2025 (*14*).

The increase in herd incidence and the recalcitrant nature of bTB to broader eradication efforts are multifaceted and underpinned by numerous risk factors, including, but not limited to: stocking density (*15*); the presence of wildlife reservoirs of *M. bovis* (e.g., the European badger – *Meles meles*) (*16*); intrapopulation genomic variation (*17*); age and sex (*18*); and the imperfect properties of current bTB diagnostic tests for correctly classifying *M. bovis*-infected animals (i.e., low sensitivity), which enables the disease to remain within cattle herds (*19*).

There are currently two diagnostic tests used for national bTB eradication programmes in Britain and Ireland: 1) the *in vivo* field-based single intradermal comparative tuberculin test (SICTT) used in isolation or in conjunction with; 2) an ancillary *in vitro* enzyme-linked immunosorbent assay (ELISA)-based interferon-γ (IFN-γ) release assay (IGRA) test to increase the sensitivity of the diagnosis by detecting infected animals as early as 14 days post-infection (dpi) (*20, 21*). Exploiting data from a range of studies, detailed meta-analyses have yielded low sensitivity estimates of the SICTT under field conditions, ranging from 0.50 (95% posterior credible interval (Crl): γ = 95%, 0.26 ≤ μ ≤ 0.78) based on data from several different countries (*22*), 0.57 (95% Crl: γ = 95%, 0.53 ≤ μ ≤ 0.63) in Ireland (*23*), to 0.66 (95% confidence interval (CI); 0.52 − 0.8) in Great Britain (*24*). In contrast, the IGRA test is considered more sensitive than the SICTT, with estimates ranging from 0.63 to 0.88 in Ireland (*20, 23*), 0.67 (95% Crl: γ = 95%, 0.49 ≤ μ ≤ 0.82) based on a statistical meta-analysis of data from different countries (*22*), and 0.93 (95% Crl: γ = 95%, 0.90 ≤ μ ≤ 0.96) in Northern Ireland (*25*). Conversely, the SICTT is considered more specific than the IGRA, expected to misclassify one bTB-negative (bTB−) animal as bTB-positive (bTB+) for every 5000 tests conducted and, as such, is selected as the primary population screening test for bTB (*26*).

Previous work in humans has shown that peripheral blood (PB) transcriptional biomarkers identified using microarray or RNA sequencing (RNA-seq) technologies can accurately discriminate between active hTB cases and asymptomatic latent tuberculosis infection patients or other infectious diseases, and also between individuals with active hTB and healthy controls (*27–29*). In cattle, PB immune responses during bTB have been shown to reflect those at the site of infection (*30*), and recent work has shown that there is substantial overlap of differentially expressed genes (DEGs) in bovine alveolar macrophages (AMs) experimentally infected with *M. bovis* and PB RNA-seq data from cattle naturally infected with *M. bovis* (*31, 32*). However, few studies have attempted to characterise PB transcriptional biomarkers capable of discriminating between bTB− and bTB+ cattle with appropriate discovery (i.e., training) and validation (i.e., testing) cohorts. For example, using microarray data, a set of 15 DEGs derived from peripheral blood mononuclear cells (PBMCs) was proposed as being capable of indicating *M. bovis* infection status (*33*), but this failed to replicate in an analysis of peripheral blood leukocytes (PBLs) from cattle naturally infected with *M. bovis* (*34*). A recent time series study of cattle experimentally infected with *M. bovis* revealed a set of 17 DEGs exhibiting increased expression that were consistently differentially expressed (DE) in PB at +1 week post-infection (wpi), +2 wpi, +6 wpi, +10 wpi, and +12 wpi, which showed striking similarity to PBL RNA-seq and microarray results in an independent cohort of animals (*34–36*). However, only three of these genes were DE in an analysis of substantially large numbers of bTB− and bTB+ cattle, and only one gene (*FOSB*) showed increased expression (*31*).

Given the zoonotic threat of *M. bovis*, the global economic cost of bTB, the increasing prevalence of the disease in Ireland and the persistence of infection in England and Wales (*37*), there is an urgent need to develop new diagnostics that can accurately discriminate between bTB− and bTB+ cattle with a high degree of specificity and sensitivity. Therefore, the objectives of this study were to leverage PB transcriptome data from bTB− cattle and cattle experimentally or naturally infected with *M. bovis* to train and evaluate a range of machine learning (ML) models in terms of their ability to discriminate between bTB− and bTB+ cattle, and also cattle infected with other viral and bacterial pathogens, including bovine herpes virus 1 (BoHV-1), bovine respiratory syncytial virus (BRSV) and *Mycobacterium avium* ssp. *paratuberculosis* (**Fig. 1**).

**Fig. 1:**
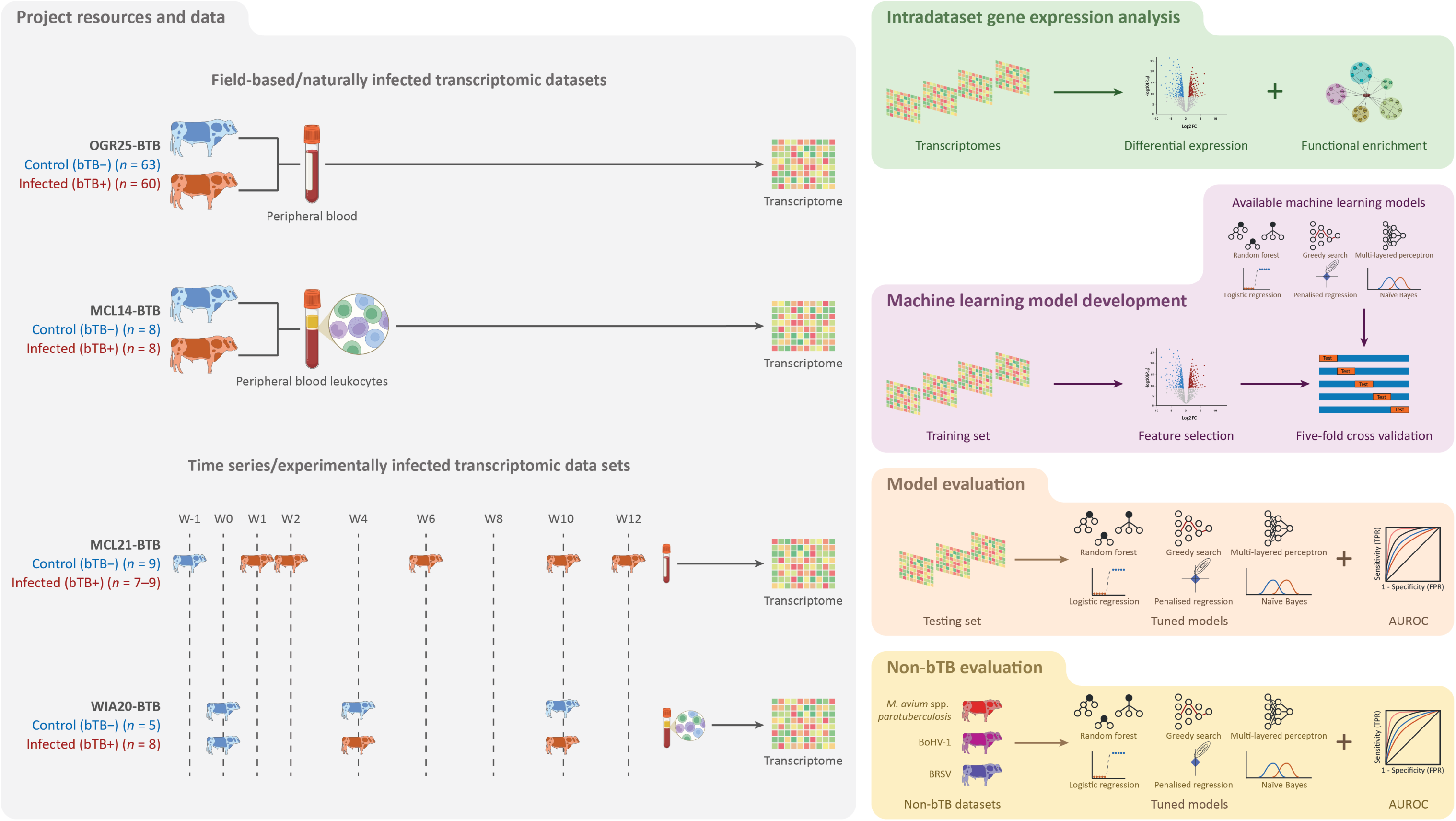
Experimental and computational workflow. Data resources for the project included peripheral blood (PB) and peripheral blood leukocyte (PBL) RNA-seq data from four studies; Two studies from (*31*) (OGR25-BTB) and (*35*) (MCL14-BTB) investigating transcriptional differences between non-infected animals (bTB−) and cattle naturally infected with *M. bovis* (bTB+), and two studies consisting of bTB− and bTB+ cattle experimentally infected with *M. bovis* and longitudinally sampled across an experimental time course from (*36*) (MCL21-BTB) and (*38*) (WIA20-BTB). Data analysis procedures included: (1) an intra-dataset differential expression (DE) and functional enrichment analysis; (2) development and tuning of machine-learning models in the training set via five-fold cross-validation using differentially expressed genes identified in the training set (see “Methods”); (3) evaluation of hyperparameter-tuned models in the testing set and assessing performance by calculating the area under the receiver operating characteristic curve (AUROC), sensitivity and specificity, respectively; and (4) evaluation of hyperparameter-tuned models in non-bTB datasets comprised of non-infected animals and cattle infected with *M. avium* spp. *paratuberculosis* (MAP), bovine herpes virus (BoHV−1), or bovine respiratory syncytial virus (BRSV), respectively.

## Results

### Intra-dataset differential expression analysis highlights substantial heterogeneity present among bTB+ animals

We performed a case-control differential gene expression analysis (DEA) to examine transcriptional changes in PB and PBL induced by *M. bovis* infection and bTB disease. To do this, we compared PB and PBL RNA-seq data from a total of *n* = 230 animals, either naturally or experimentally infected with *M. bovis* (bTB+), and control non-infected cattle (bTB−), enrolled in four studies conducted in Ireland, the UK, and the US. These included cattle naturally infected with *M. bovis* from (*31*) (OGR25-BTB) and (*35*) (MCL14-BTB) in addition to cattle experimentally infected with *M. bovis* from (*36*) (MCL21-BTB) and (*38*) (WIA20-BTB) that were longitudinally sampled across an experimental infection time course (**Table 1**, **Fig. 1**).

**Table 1:** Overview of datasets used in this manuscript.

| Study | Infection acquisition | Disease studied | Abbreviation | Sample size | Sampling points | Sampling time points weeks post-infection (wpi) | Total number of RNA-seq samples | Tissue type |
| --- | --- | --- | --- | --- | --- | --- | --- | --- |
| (31) | Natural | bTB<br>(*M. bovis*) | OGR25-BTB | $n = 123$<br>(63 bTB–, 60 bTB+) | 1 | - | 123 | PB |
| (35) | Natural | bTB<br>(*M. bovis*) | MCL14-BTB | $n = 16$<br>(8 bTB–, 8 bTB+) | 1 | - | 16 | PBL |
| (36) | Experimental | bTB<br>(*M. bovis*) | MCL21-BTB | $n = 7-9$<br>(paired design) | 5 | –1 wpi, +1 wpi, +2 wpi,<br>+6 wpi, +10 wpi, +12 wpi | 52 | PB |
| (38) | Experimental | bTB<br>(*M. bovis*) | WIA20-BTB | $n = 13$<br>(5 bTB–, 8 bTB+) | 3 | 0 wpi, +4 wpi, +10 wpi | 39 | PBL |
| (39) | Natural | Johne’s Disease<br>(MAP) | ALO19-MAP | $n = 14$<br>(3 MAP–, 11 MAP+) | 1 | - | 14 | PB |
| (40) | Experimental | BRD<br>(BRSV) | JOH21-BRD | $n = 18$<br>(6 BRSV–, 12 BRSV+) | 1 | - | 18 | PB |
| (41) | Experimental | BRD<br>(BoHV-1) | ODO23-BRD | $n = 18$<br>(6 BoHV–1–, 12 BoHV–1+) | 1 | - | 18 | PB |

bTB, bovine tuberculosis; MAP, *M. avium* ssp. *paratuberculosis*; BRD, bovine respiratory disease; BRSV, bovine respiratory syncytial virus; BoHV–1, bovine herpes virus 1; PB, peripheral blood; PBL, peripheral blood leukocytes.

We first analysed each of the four datasets individually to characterise differentially expressed genes (DEGs) that were consistent between studies. Principal component analysis (PCA) within each dataset showed variable separation of bTB+ from bTB− animals, likely reflecting differences in disease stage (e.g., acute versus chronic), experimental infection dose, age, or genetic variation due to breed composition differences. With respect to the naturally infected cattle from the MCL14-BTB dataset, we observed a distinct separation between bTB+ and bTB− animals, with PC1 accounting for 49% of the total variance explained in the PBL transcriptomes (**Fig. S1a**). Conversely, in the OGR25-BTB dataset, we observed indistinct separation of the bTB+ and bTB− cattle, with PC1 accounting for 12% of the total variance (**Fig. S1b**). For the PB MCL21-BTB time series study, after experimental infection with *M. bovis*, we observed weak separation of bTB+ and bTB− cattle at +2 wpi; however, a clear division between the two cohorts was evident at +10 wpi, which diminished markedly by +12 wpi (**Fig. S2**). Similarly, for the PBL WIA20-BTB, the distinction between bTB+ and bTB− cattle was clear at +4 wpi but was substantially reduced at +10 wpi (**Fig. S3**).

Within each dataset, we subsequently conducted a DEA between the bTB+ and bTB− cattle, incorporating the top two PCs derived from the RNA-seq-called SNP variants within each dataset to account for functional genetic variation due to breed/population differences among the animals (**Fig. S4**). We observed that for variants called across all datasets, and which were present in the (*31*) imputed and filtered WGS dataset, broad population structure was captured: the coordinates of eigenvector 1 and 2 corresponding to PC1 and PC2, respectively, for the *n* = 123 animals described by (*31*) displayed a significant and positive Spearman correlation (*ρ*) with the eigenvector coordinates for the same set of animals derived from pruned high-density SNP-array data (**Fig. S5a** and **Fig. S5b**).

In total, 13,897 genes were consistently expressed across all datasets, and we identified 4,533 DEGs in the MCL14-BTB dataset, with 2,223 and 2,310 genes displaying increased and decreased expression, respectively, in the bTB+ group compared to the bTB− cohort. For the OGR25-BTB dataset, we characterised 3,139 DEGs, of which 1,969 showed increased expression, whereas 1,170 showed decreased expression in the bTB+ set compared to the bTB− group. For the MCL21-BTB time series experiment, we identified 2,769 DEGs across all time points (2,402 unique DEGs). Of these, 19, 55, 226, 1,095, and 48 displayed increased expression in the bTB+ group compared to the bTB− cohort (−1 wpi) at +1 wpi, +2 wpi, +6 wpi, +10 wpi, and +12 wpi, respectively. Conversely, 3, 2, 93, 1,212, and 16 displayed decreased expression in the bTB+ group compared to the bTB− animals at +1 wpi, +2 wpi, +6 wpi, +10 wpi, and +12 wpi, respectively. For the WIA20-BTB dataset, across all time points, we identified 3,124 DEGs (2,434 unique DEGs). Of these, 1,246 and 623 displayed increased and decreased expression, respectively, in the bTB+ group at +4 wpi versus the bTB− group. At 10 wpi in the bTB+ group, 627 and 628 DEGs displayed increased and decreased expression, respectively, compared to the bTB− group (**Fig. 2A**, **Table S1**).

**Fig. 2:**
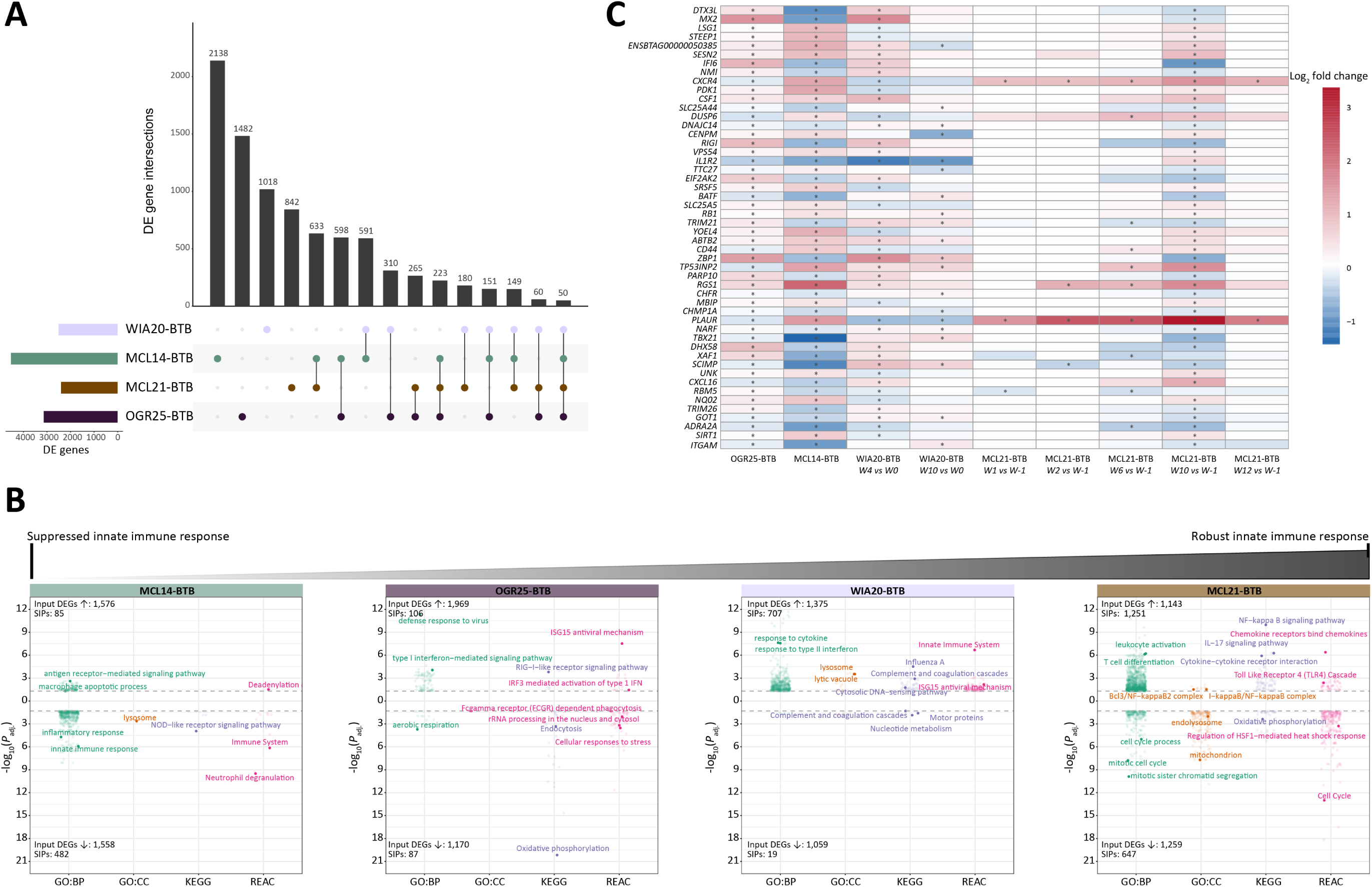
Intra-dataset differential expression and functional enrichment analysis. (A) Upset plot illustrating the number of significantly (*P*_adj._ < 0.05) differentially expressed genes (DEGs) identified in the four bovine tube culosis (bTB) datasets and the overlap of these DEGs among the studies. (B) Bi-directional jitter plots for significantly impact pathways (SIPs) perturbed by DEGs exhibiting increased and separately, decreased expression in each dataset, respectively for four databases: (1) gene ontology (GO) biological processes (GO:BP); (2) GO cellular component (GO:CC); (3) Kyoto Encyclopaedia of Genes and Genomes (KEGG); and (4) Reactome (REAC). Dashed dotted lines indicate the −log_10_*P*_adj._ threshold of 0.05 for characterising a SIP. The total number of input DEGs and identified SIPs are also detailed for each analysis of increased and decreased DEGs in each dataset, respectively. The jitter plots are arranged from left to right according to their documented level of innate immune response pathway activation. (**C**) Heatmap showing the log_2_ fold-change (LFC) in effect size estimates between control non-infected cattle and animals infected with *M. bovis* for 50 significant DEGs between bTB− and bTB+ cattle identified in all naturally infected datasets and in at least one time point in each of the time course datasets, respectively. Red colours indicate increased expression, and blue colours indicate decreased expression in the bTB+ group relative to the bTB− group. The * sign indicates that the LFC value was significantly greater than or less than 0 in the corresponding comparison between bTB+ and bTB− cattle.

We used g:Profiler to evaluate the biological pathways perturbed by DEGs, separately for those exhibiting increased and decreased expression within each dataset. We observed that these DEGs significantly impacted (*P*_adj._ < 0.05) multiple immune response pathways, illustrating the broad spectrum of the host response to *M. bovis* infection, which ranged from a suppressed innate immune profile in the MCL14-BTB to one that was highly active in the MCL21-BTB dataset (**Fig. 2B**, **Table S2**). In the MCL14-BTB dataset, many of the decreased DEGs were in significantly impacted pathways (SIPs) such as *Inflammatory response*, *NOD-like receptor signalling pathway*, and *Immune system*, likely reflecting suppression of these key innate immune pathways in response to chronic *M. bovis* infection. For the OGR25-BTB dataset, the results were more heterogeneous, likely reflecting the diverse bTB disease states within this study cohort. For example, SIPs perturbed by the increased DEGs included classical biological processes perturbed during *M. bovis* infection (*32*), such as *RIG-I-like receptor signaling*, *ISG15 antiviral mechanism*, and *positive regulation of T-helper 2 cell cytokine production*. Conversely, the decreased DEGs impacted the *Cellular responses to stress* and *FCgamma receptor dependent phagocytosis* signalling pathways. For the time series experiments, many of the increased DEGs were involved in biological processes reflecting an acute and robust innate immune response to *M. bovis* infection. These pathways included *Response to type II interferon*, *Cytosolic DNA-sensing pathway*, *IL-17 signaling pathway*, *TLR4 cascade*, *Leukocyte activation*, and *NF−κB signaling* (**Table S2**)

The overlap of DEGs identified among the studies is shown in **Fig. 2A**. In total, we identified 50 DEGs in the naturally infected *M. bovis* datasets that were also significantly DE in at least one time point of the time series experiments (**Fig. 2A**, **Fig. 2C**). Analysis of the log_2_ fold-change (LFC) values of the DEGs for each contrast revealed a heterogeneous pattern (**Fig. 2C**); For example, the *PLAUR* gene was significantly DE across all nine contrasts and had a positive LFC value in the MCL14-BTB and MCL21-BTB datasets but a negative LFC value in the OGR25-BTB and WIA20-BTB datasets, respectively. Conflicting results were also observed for a range of other immune related genes including *CXCL16*, *IFI6*, *RIGI*, and *CXCR4*. However, albeit not significant across all comparisons, six genes (*SESN2, VPS54*, *RB1*, *CSF1*, *ABTB2*, and *RGS1*) possessed a positive LFC value in each contrast whereas one gene (*ADRA2A*) displayed decreased expression in bTB+ animals for each contrast.

### Consolidation of bTB datasets identifies transcriptional biomarkers that accurately discriminate between bTB+ and bTB− animals

We consolidated the four bTB datasets and split all samples (*n* = 230) randomly into a training set (70%; *n* = 162; *n* = 69 bTB−, *n* = 93 bTB+) and a testing set (30%; *n* = 68; *n* = 34 bTB−, *n* = 34 bTB+). The animal IDs and associated metadata of samples split into the training and testing sets are described in **Tables S3** and **S4**. Consolidation of the four bTB datasets revealed the presence of a strong batch effect, whereby samples clustered in a study-specific manner, a technical artefact that was preserved in both the training and testing sets (**Fig. S6**). Within the training set, a DEA was conducted between bTB+ and bTB− cattle accounting for age, study/batch, and the top two genotype PCs inferred from SNPs called from the RNA-seq data, which were intersected and merged with the filtered imputed WGS data from (*31*) (**Table S3**, **Fig. S7**). From this analysis, a total of 1,635 DEGs were identified: 949 genes exhibited increased expression and 686 displayed decreased expression in the bTB+ group relative to the bTB− cohort. Of these, 730 increased and 581 decreased DEGs were retained for the machine learning analysis, as they had a median-of-ratios expression value > 100 across samples in the training set (**Fig. 3A**, **Table S5**). Significantly impacted pathways (*P*_adj._ < 0.05) enriched for the retained DEGs exhibiting increased expression in the bTB+ group included: *Innate immune response*, *Regulation of tumor necrosis factor production*, the *Interferon-alpha production* GO:BP terms; the *NOD-like receptor signaling pathway* KEGG term; and the *Interleukin-21 signaling* Reactome term. For the retained DEGs displaying decreased expression, SIPs enriched for these genes included: the *Regulation of macroautophagy* and *Regulation of interleukin-1-mediated signaling pathway* GO:BP terms, and the *Mitotic Spindle Checkpoint* Reactome term (**Fig. 3B**, **Table S6**).

**Fig. 3:**
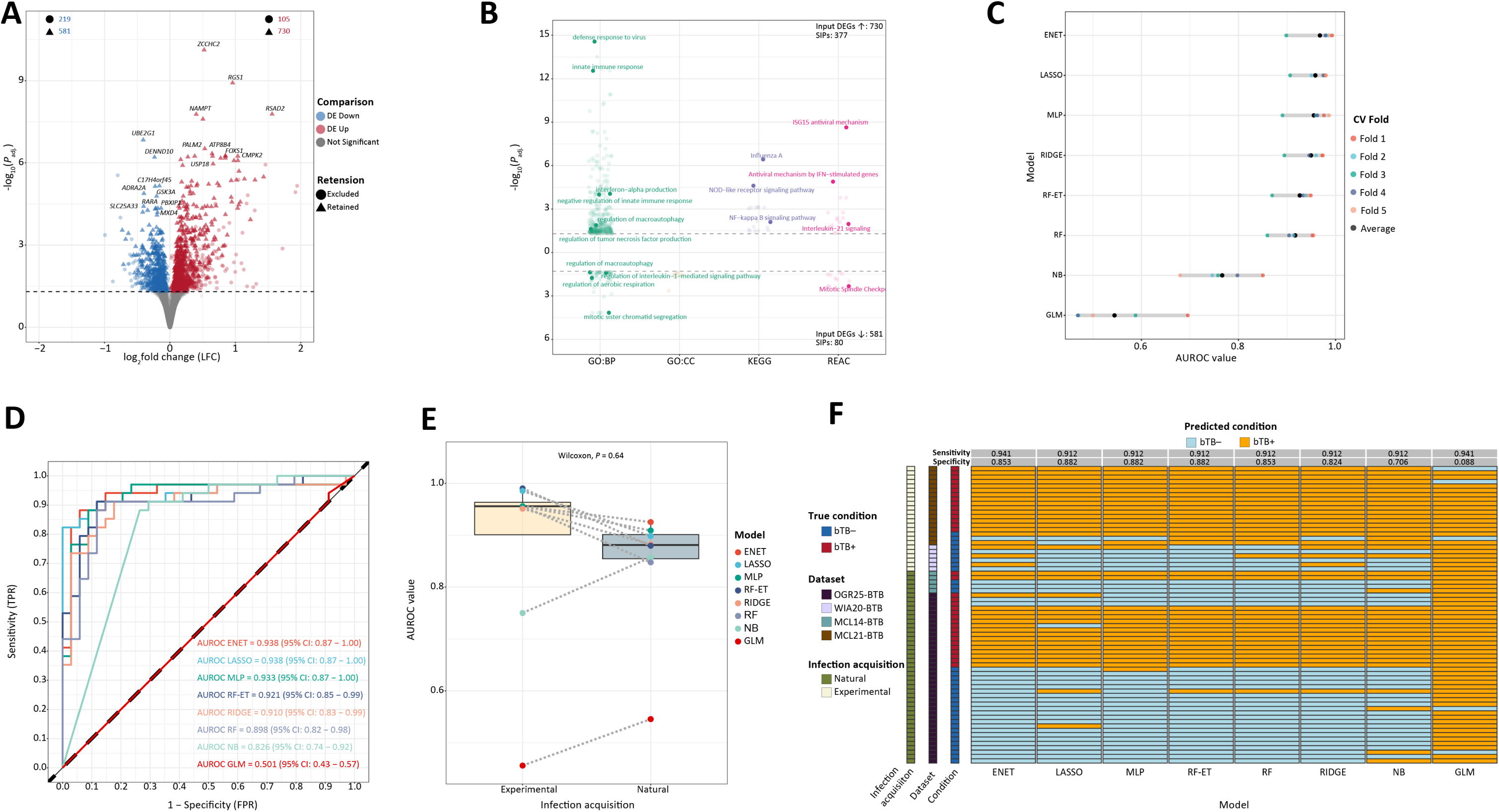
Machine learning (ML) model training and evaluation. (**A**) Volcano plot illustrating significantly differentially expressed genes (DEGs) identified in the training set for the bTB+ (*n* = 93) vs bTB− (*n* = 69) contrast with thresholds determined by FDR-*P*_adj._ < 0.05 and an absolute log_2_ fold-change (LFC) > 0. Genes are coloured to indicate increased (red) or decreased (blue) expression in the bTB+ group relative to the bTB− group, respectively, and triangular datapoints indicate the genes retained for subsequent ML analysis with the criterion of mean median-of-ratios normalised expression count >100 in the training dataset. (**B**) Bi-directional jitter plot for significantly impacted pathways (SIPs) perturbed by DEGs exhibiting increased and separately, decreased expression in the training set for four databases: (1) gene ontology (GO) biological processes (GO:BP); (2) GO cellular component (GO:CC); (3) Kyoto Encyclopaedia of Genes and Genomes (KEGG); and (4) Reactome (REAC). Dashed dotted lines indicate the −log_10_*P*_adj._ threshold of 0.05 for characterising a SIP. The total number of input DEGs and identified SIPs are also detailed for each analysis of increased and decreased genes in each dataset, respectively. (**C**) Area under the receiver operating characteristic curve (AUROC) values for eight hyperparameter-tuned ML models in each of the five folds of the training set. (**D**) An AUROC depicting the performance of each of the hyperparameter-tuned models in the testing set. (**E**) Boxplot showing the distribution of AUROC estimates of animals in the testing dataset depending on whether the dataset comprised of animals experimentally or naturally infected with *M. bovis*. (**F**) Heatmap showing the predicted class of animals in the testing set when selecting an optimum threshold to maximise the sensitivity (> 0.90), where possible, of each classifier.

We trained and performed hyperparameter tuning of eight models (unpenalised generalised logistic model [GLM], logistic regression with either a lasso [LASSO], ridge [RIDGE] or elastic net [ENET] penalisation, respectively, random forest [RF] and random forest specifying the extra trees algorithm [RF-ET], naïve Bayes [NB], and a multi-layered perceptron [MLP]) in the training set using five-fold cross-validation. The selected hyperparameters for each model are described in **Table S7**. The performance of each tuned model across the five folds is illustrated in **Fig. 3C** and described in **Table S8**. The best performing model was the ENET model, which achieved an average AUROC of 0.968 (range: 0.898–0.993). Other models that performed almost as well included the LASSO (0.958, [0.906–0.981]), the MLP (0.955, [0.891–0.987]), the RIDGE (0.949, [0.895–0.974]), the RF-ET (0.926, [0.869–0.949]), and the RF (0.917, [0.859–0.955]) classifiers. The NB classifier produced a substantially lower average AUROC value of 0.766 (0.679–0.850). The poorest-performing model in the training set was the GLM, which gave an average AUROC value across the five folds of 0.545 (0.469–0.696) (**Fig. 3C**, **Table S8**).

Evaluation of the hyperparameter-tuned models in the testing set illustrated that all the models generalised well, achieving comparable performance to that observed in the training set (**Fig. 3D**). The best-performing models were the ENET and the LASSO models, which both achieved an AUROC of 0.938 (95% confidence interval (CI): 0.87–1.00). These models were followed closely by the MLP (0.933, 95% CI: 0.87–1.00), RF-ET (0.921, 95% CI: 0.85–0.99), RIDGE (0.910, 95% CI: 0.83–0.99), and RF (0.898, 95% CI: 0.82–0.98) models, respectively. The NB model, similar to observations in the training set, gave substantially lower performance in the testing set, obtaining an AUROC of 0.826 (95% CI: 0.74–0.92). Again, consistent with findings from the training set analysis, the worst-performing model was the GLM model, which performed essentially like random guessing (AUROC = 0.501, 95% CI: 0.43–0.57). Taking the AUROC value as the performance metric, we observed that for all models except the NB and GLM classifiers, the ability to classify animals correctly as bTB+ or bTB−, respectively, in the experimentally infected datasets tended to be higher than for the naturally infected datasets; however, this difference was not statistically significant (Wilcoxon paired rank-sum test; *P* = 0.64; **Fig. 3E**).

Current bTB diagnostics lack sensitivity. We therefore exploited the ROC curves depicted in **Fig. 3D** and for each model, selected an optimum probability threshold (*θ*) for classifying an animal as bTB+ that yielded a sensitivity value > 90% where possible, while also balancing specificity estimates (**Table S9**). At these selected *θ* values, the models that obtained the joint highest sensitivity estimates were the GLM and ENET classifiers, each achieving a sensitivity value of 0.941; however, the GLM model misclassified a high proportion of bTB− animals as bTB+ (31 out of 34 bTB− animals), yielding a specificity estimate of 0.088 (**Fig. 3F**, **Table S10**). Overall, the models that achieved the optimal balance between sensitivity and specificity were the ENET classifier (sensitivity = 0.941, specificity = 0.853), followed closely by the LASSO, MLP, and RF-ET models (sensitivity = 0.912, specificity = 0.882). Similar sensitivity estimates were observed for the RF and RIDGE models, with reduced specificity estimates (0.853 and 0.824, respectively). The NB model obtained comparable sensitivity estimates to the LASSO, MLP, RF-ET, RF, and RIDGE models (0.912), but had substantially lower specificity (0.706). The ENET model incorrectly classified two out of 34 bTB+ animals (5.88%) as bTB−, with both animals being derived from the OGR25-BTB dataset. All models, apart from the GLM classifier, correctly classified the animals experimentally infected with *M. bovis* as bTB+ from the MCL21-BTB time series dataset, which included 3 bTB+ animals sampled at +1 wpi and +2 wpi (**Fig. 3F**, **Table S10**). This is an important result because the current IGRA bTB diagnostic test has a minimum +2 wpi chronological detection limit, because an adaptive immune response to mycobacterial antigens is required for the assay to function (*21*).

### A greedy forward search algorithm identifies robust discriminatory gene sets in the training set with poor generalisability in the testing set

Previous studies have shown the utility of employing a greedy forward search (GFS) algorithm for identifying a small number of discriminatory genes between non-infected individuals and individuals diagnosed with a range of diseases, including sepsis (*42*), dengue fever (*43*), and hTB using both microarray (*28*) and RNA-seq data (*29*), respectively. We therefore implemented the GFS algorithm (see “Methods”) on the entire set of retained DEGs (1,311 genes; **Fig. 3A**). This analysis identified a parsimonious set of 13 genes (*SATB1*, *ALAS1*, *NR4A2*, *SMG8*, *ELP2*, *DCAF5*, *ZNF75D*, *PRICKLE3*, *MYB*, *BCDIN3D*, *TEX2*, *MAD2L2*, and *KATNBL1*; 13-gene set, **Fig. 4A**), which achieved an average AUROC across the five folds of 0.996 (range: 0.990–1.000) (**Fig. 4A**, **Fig. 4B**, **Table S11**). We subsequently reimplemented the GFS strategy on the set of retained DEGs without the genes identified in the first initialisation. This identified a set of 17 genes (*PTK2*, *TNFRSF13B*, *UBE2G1*, *SENP3*, *UQCC2*, *TAP2*, *MON1A*, *NFE2L3*, *GTSE1*, *TRIQK*, *ADPGK*, *GDPG1*, *CCR5*, *ZNF628*, *NABP1*, *GEMIN5*, and *GDAP2*; 17-gene set, **Fig. 4A**), which achieved near perfect discrimination across the five folds in the training set, with an average AUROC across the five folds of 0.999 (range: 0.993–1.000) (**Fig 4A**, **Fig. 4B**, **Table S12**). We also evaluated a combination of the 13-gene and 17-gene sets (30-gene set), respectively, which yielded an average AUROC across the five folds in the training set of 0.986 (range: 0.975–1.000) (**Table S13**).

**Fig. 4:**
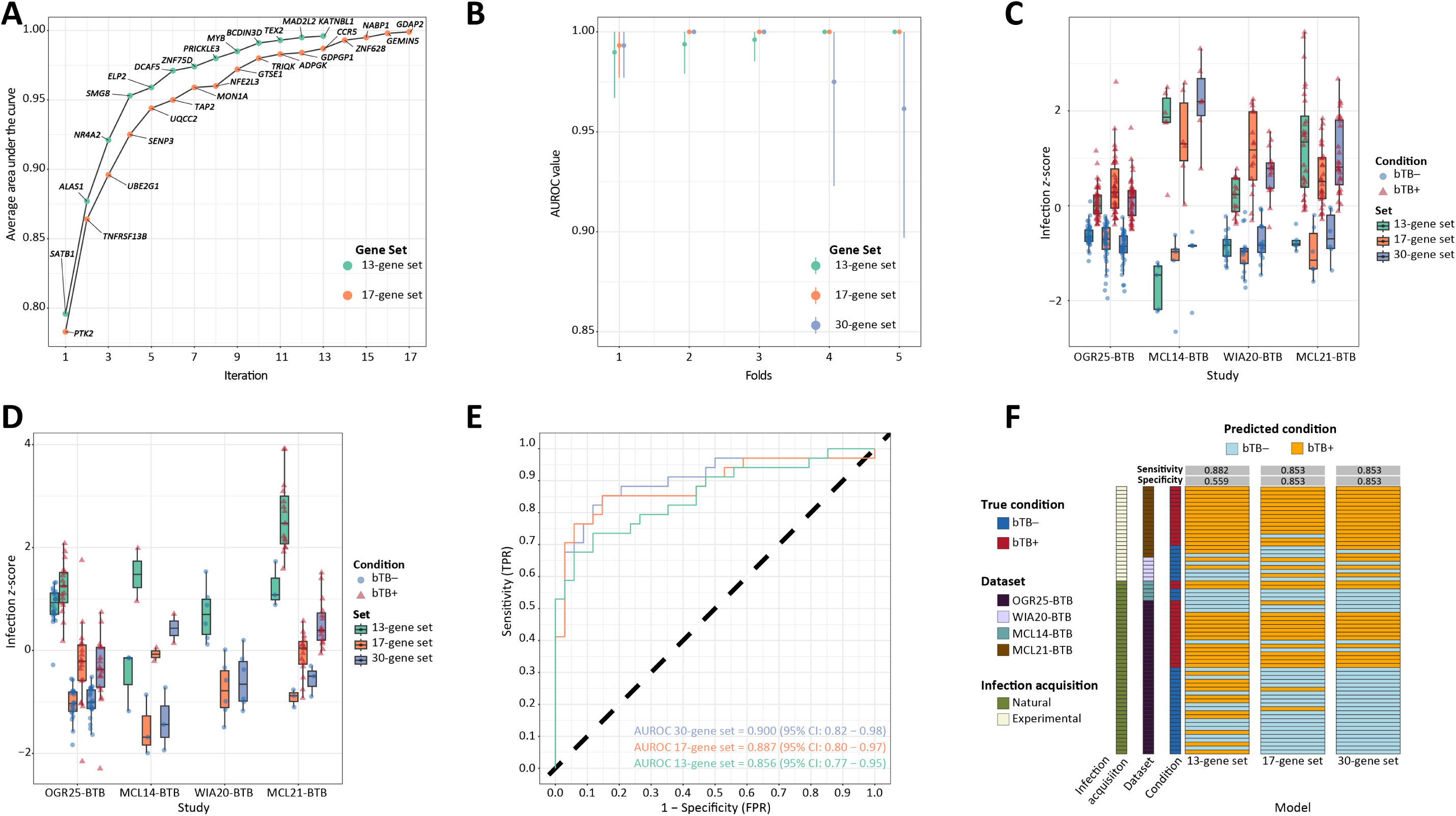
Greedy-forward search strategy evaluation in training and testing sets. (**A**) Line plot showing the average area under the receiver operating characteristic curve (AUROC) across the five folds in the training set for each combination of genes identified in the first pass or second pass of the greedy-forward search algorithm, respectively. (**B**) Line plot showing the 95% confidence interval of AUROC estimates for the 13-gene set, 17-gene set and combined 30-gene set in each of the five cross-validation folds in the training set. (**C**) Boxplots showing the distribution of the scaled infection *z*-score derived from each of the three gene sets for all bTB− (blue; circle) and bTB+ (red; triangle) animals in the training set, separated by study of origin. (**D**) Boxplots showing the distribution of the scaled infection *z*-score derived from each of the three gene sets for all bTB− (blue) and bTB+ (red) animals in the testing set, separated by study of origin. (**E**) An AUROC depicting the performance of each of the three gene sets in the testing set. (**F**) Heatmap showing the predicted class of animals in the testing set when selecting an optimum threshold to maximise the sensitivity (> 0.85) for each of the three gene sets.

Evaluation of the infection *z*-scores (see “Methods”) inferred from each of the gene sets across all four cohorts in the training dataset revealed significant differences between bTB+ and bTB− cattle, with all intra-dataset-condition pairs showing a significant difference in the inferred infection *z-*score (Wilcoxon rank-sum test, *P*_adj._ < 0.05) (**Fig. 4C, Table S14)**.

For each gene set, we next inferred the infection *z-*score of bTB+ and bTB− animals in the testing set and observed significant differences in the inferred bTB scores for these conditions in the MCL21-BTB and OGR25-BTB datasets (Wilcoxon rank-sum test, *P*_adj._ < 0.05); however, this difference was not statistically significant in the MCL14-BTB dataset (*P*_adj._ = 0.20) owing to the small sample size (*n* = 2 bTB+, *n* = 3 bTB−) (**Fig. 4D**, **Table S15**). We were unable to compare the bTB score between bTB+ and bTB− animals in the WIA20-BTB cohort in the testing set because the original train/test split yielded no bTB+ animals from this cohort in the testing set. Evaluation of the predictive capabilities of the three gene sets revealed significantly lower AUROC estimates in comparison to the training set with the 13-gene set, the 17-gene set and the 30-gene set achieving AUROC values of 0.856 (95% CI: 0.77−0.95), 0.887 (95% CI: 0.80−0.97), and 0.900 (95% CI: 0.82−0.98), respectively (**Fig. 4E**).

Following the same approach used for the analysis of the other ML algorithms (**Fig. 3F**), for each gene set, we selected the threshold that yielded a sensitivity value > 0.85 and that also maximised the specificity (**Table S16**). We selected a lower sensitivity value in this analysis because selecting a sensitivity > 0.9 for all three gene sets yielded very low specificity estimates (< 0.65 for all sets; **Table S16**). Using these selected threshold values, the 13-gene set achieved a sensitivity of 0.882 and a specificity of 0.559, with a large number of bTB− cattle misclassified as bTB+ (15 out of 34 bTB− animals). The 17-gene and 30-gene set signatures both achieved balanced sensitivity and specificity estimates of 0.853 for each metric (**Fig. 4F**, **Table S17**). Similar to the other ML models, at these thresholds, the 13-gene and 30-gene sets correctly classified the animals experimentally infected with *M. bovis* at +1 wpi and +2 wpi, respectively, in the MCL21-BTB dataset as bTB+.

### Evaluation of comparator pathogen datasets demonstrates the robustness and specificity of classifiers and gene sets

Given that our bTB ML models and bTB gene sets showed high discriminatory power between bTB− and bTB+ cattle, respectively, we next wanted to evaluate the capacity of these classifiers to differentiate *M. bovis*-infected cattle from cattle infected with other bacterial and viral pathogens as an indicator of their specificity. To do so, we obtained RNA-seq data for two other infectious diseases caused by three pathological agents, including Johne’s disease caused by *M. avium* spp. *paratuberculosis* (MAP) (*n* = 3 MAP−, *n* = 11 MAP+) from (*39*) (ALO19-MAP) and bovine respiratory disease (BRD) caused by infection with either BoHV-1 (*n* = 6 BoHV-1−, *n* = 12 BoHV-1+) from (*41*) (ODO23-BRD) or BRSV (*n* = 6 BRSV−, *n* = 12 BRSV+) from (*40*) (JOH21-BRD), respectively (**Table 1**, **Fig. 1**).

Within each dataset, we first evaluated the ability of the eight hyperparameter-tuned ML classifiers, trained on the bTB training data, to discriminate between control non-infected animals and cattle infected with each of the three pathogens. With regards to the MAP dataset, we observed that all eight ML models performed poorly, with the RIDGE model achieving the highest AUROC value of 0.667 and the RF-ET model achieving the lowest AUROC of 0.455, no better than random allocation; however, wide confidence interval estimates were inferred because of the small number of animals in this dataset (**Fig. 5A**). For animals infected with BoHV-1, we observed that the models displayed improved discriminative ability, achieving a maximum AUROC from the ENET model of 0.931 (95% CI: 0.79–1.00), whereas the NB and GLM models achieved the lowest AUROC of 0.50 (95% CI: 0.5−0.5) (**Fig. 5B**). For cattle experimentally infected with BRSV, we observed high discriminatory ability with the LASSO, RIDGE, RF, RF−ET, RIDGE, and ENET models, all achieving an AUROC ≥ 0.944, whereas as the GLM and NB models achieved an AUROC of 0.667 and 0.500, respectively (**Fig. 5C**).

**Fig. 5:**
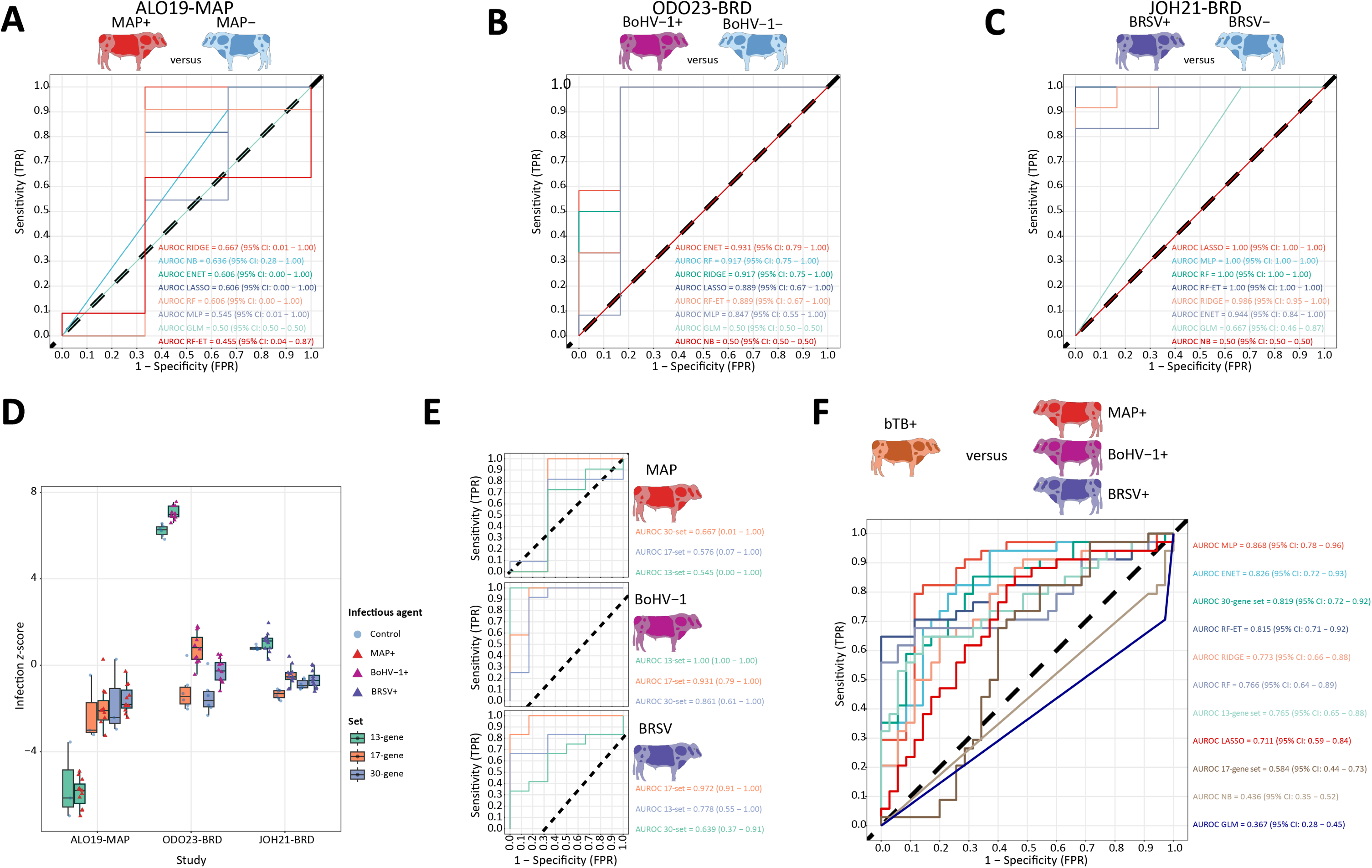
Evaluation of machine learning and greedy forward search strategies in external datasets. (**A**) An area under the receiver operating characteristic curve (AUROC) showing the ability of the hyperparameter-tuned models to discriminate between control non-infected animals and cattle infected with *M. avium* spp. *paratuberculosis* (MAP) from (*39*) (ALO19-MAP) dataset. (**B**) An AUROC showing the ability of the hyperparameter-tuned models to discriminate between control non-infected animals and cattle infected with bovine herpes virus (BoHV-1) from (*41*) (ODO23-BRD) dataset. (**C**) An AUROC showing the ability of the hyperparameter-tuned models to discriminate between control non-infected animals and cattle infected with bovine respiratory syncytial virus (BRSV) from (*40*) (JOH21-BRD) dataset. (**D**) Boxplots showing the distribution of the scaled infection *z*-score derived from each of the three gene sets for all control non-infected animals and cattle infected with either MAP, BoHV-1, or BRSV, respectively. (**E**) Three AUROCs illustrating the performance of the three gene sets in terms of discriminating between control non-infected animals and cattle infected with either MAP, BoHV−1, or BRSV, respectively. (**F**) An AUROC curve showing the discriminatory power of all hyperparameter-tuned ML models and gene sets when comparing bTB+ cattle to MAP+, BoHV-1+ and BRSV+ cattle, respectively.

For each non-bTB dataset, we next inferred an infection *z*-score for all non-infected and infected animals using the 13-gene set biosignature inferred from the first iteration of the GFS algorithm on the bTB training dataset, the 17-gene set signature from the second iteration, and the combined 30-gene set inferred from both iterations (**Fig. 4A**, **Fig. 5D**). We observed significant differences (Wilcoxon rank-sum test; *P*_adj._ < 0.05) between the inferred infection *z*-scores from all three gene set signatures for control non-infected animals and cattle infected with BoHV-1 (**Fig. 5D**, **Table S18**). Conversely, this difference was not significant between MAP+ and MAP− animals for any of the three gene set signatures (*P*_adj._ ≥ 0.586), or between BRSV+ and BRSV− animals using either the 13-or 30-gene set signatures, respectively (*P*_adj._ ≥ 0.120) (**Table S18**). However, the difference in the inferred infection *z*-score between BRSV+ and BRSV− animals was significant based on the 17-gene set signature (*P*_adj._ = 1.9 × 10^−3^) (**Fig. 5D**, **Table S18**).

We next inferred the discriminatory ability of these gene sets between control non-infected and infected animals in each non-bTB dataset using ROC curves **(Fig. 5E**). We observed that the three gene sets performed poorly at discriminating between MAP+ and MAP− animals, achieving an AUROC ≤ 0.667; however, these estimates were accompanied by wide confidence intervals. For the BoHV-1 dataset, similarly to the observations from the ML models, we observed better discriminatory capacity between BoHV-1− and BoHV-1+ animals with the 30-, 17-, and 13-gene set signatures achieving AUROC estimates of 0.861, 0.931, and 1.00 (perfect discrimination), respectively (**Fig. 5E**). With respect to the BRSV dataset, we observed high discriminatory power for the 17-gene set signature, which achieved an AUROC of 0.972 (95% CI: 0.91–1.00). Conversely, the 13- and 30-gene set signatures performed poorly at distinguishing between BRSV− and BRSV+ cattle, yielding AUROC estimates of 0.778 and 0.639, respectively (**Fig. 5E**).

We lastly evaluated the capacity of all eight pretrained and hyperparameter-tuned ML models and the three inferred GFS gene sets to discriminate between bTB+ animals from the testing set, and animals infected with MAP, BoHV-1, or BRSV, respectively. We observed variable performance among ML models and gene sets in their ability to differentiate between bTB+ animals and animals infected with the other infectious agents (**Fig. 5F**). The best-performing model/gene set was the MLP model. This model yielded an AUROC of 0.868 (95% CI: 0.78–0.96), indicating that it is highly specific in differentiating between animals with bTB and animals with other bacterial and viral infections. This was closely followed by the ENET model (the best performing model for distinguishing between bTB+ and bTB− cattle; **Fig. 3C**), which achieved an AUROC of 0.826 (95% CI: 0.72–0.93) (**Fig. 5F**). The 30- and 13-gene set signatures, and the RF-ET, RIDGE, and RF models, performed comparably well, obtaining AUROC estimates ranging from 0.765−0.819. We subsequently observed a drop-off in performance for the LASSO and 17-gene set signature, which achieved AUROC values of 0.711 and 0.584, respectively. The NB and unpenalised GLM algorithms performed extremely poorly, achieving AUROC estimates < 0.5 (**Fig. 5F**).

## Discussion

We present a comprehensive, systematic evaluation and integration of PB and PBL RNA-seq data from cattle naturally or experimentally infected with *M. bovis* (bTB+) and from control, non-infected animals (bTB−). It is well established that the peripheral immune responses of cattle infected with *M. bovis* reflect those at the site of infection (*30*). In this regard, we have shown that the peripheral transcriptomic response following *M. bovis* infection can exist on a spectrum, ranging from a weak innate immune response in animals naturally infected with *M. bovis* to an immune signature that is biased towards intensive activation of canonical innate immune response pathways in animals shortly after being experimentally infected with *M. bovis*. By consolidating multiple bTB RNA-seq datasets and using eight different ML algorithms and a GFS strategy, we observed that accurate discrimination between bTB+ and bTB− cattle can be achieved with high sensitivity and specificity, and that the results generalise well to unseen test data. We also showed that a 30-gene signature, identified using a greedy-forward search algorithm, and an ENET classifier are robust at distinguishing between bTB+ cattle and animals infected with MAP, BoHV−1, or BRSV.

Analysis of RNA-seq data from bTB+ and bTB− cattle within each bTB dataset identified thousands of DEGs **(Fig. 2A**), indicating that the peripheral transcriptome is substantially perturbed following *M. bovis* infection—an observation that is consistent with previous work focused on bovine alveolar macrophages (bAMs) experimentally infected with *M. bovis* and *M. tuberculosis* (*32, 44, 45*). However, we observed that 35.1–47.2% of DEGs identified in each bTB dataset were cohort-specific, suggesting that analysis of varying bTB disease states (e.g., acute versus chronic) and *M. bovis* infectious dose, which may differ between naturally and experimentally infected animals, can yield distinct transcriptional patterns. In this regard, the DEGs that displayed decreased expression in bTB+ cattle relative to the bTB− cattle in the MCL14-BTB dataset, were enriched in pathways that are typically associated with the initial host-pathogen interaction, as reflected, for example, in the proinflammatory responses observed in bAMs infected with *M. bovis* (*32, 44, 45*) (**Fig. 2C**). Our results for bTB+ cattle naturally infected with *M. bovis* are consistent with a previous microarray study that also detected suppression of key innate immune genes in naturally infected cattle (*33*). Conversely, for the WIA20-BTB and MCL21-BTB datasets, which encompassed animals that were recently experimentally infected with *M. bovis*, the DEGs that displayed increased expression were significantly enriched in SIP sets that were dominated by interferon-mediated signalling pathways, which is consistent with an acute phase of infection that has been reported in PB data for hTB (*27, 46*).

Our analysis of SIPs enriched for DEGs with increased or decreased expression in the bTB+ cohort compared to the bTB− cohort revealed conflicting gene expression patterns in the OGR25-BTB dataset. Many of the SIPs enriched for genes showing increased expression following *M. bovis* infection were classical innate immune response and cytosolic sensing pathways, including the *RIG-I-like receptor signalling pathway*, the *Type 1 interferon-mediated signalling pathway*, and the *ISG15 antiviral mechanism* Reactome term, all of which play key roles in the host response to mycobacterial challenge (*47–49*). On the other hand, the *Chemokine signaling*, the *FCgamma receptor dependent phagocytosis* and *Cellular responses to stress* pathways were enriched for DEGs exhibiting decreased expression in the bTB+ cohort (**Fig. 2C; Table S2**), indicating suppression of these pathways. Collectively, these findings suggest that the animals in the OGR25-BTB dataset are clinically heterogeneous, and this variation is reflected in the PB transcriptome, which is unsurprising given the large number of bTB+ animals (*n* = 60) and the lack of information on the chronology of *M. bovis* infection.

In total, across all four bTB datasets, given the observed transcriptional heterogeneity, we characterised only 50 shared DEGs, the majority of which displayed substantial inter-dataset differences in terms of inferred effect sizes (LFC values) between bTB− and bTB+ cattle, respectively (**Fig. 2B**). These findings highlight the ineffectiveness of solely relying on DEGs identified within and across different bTB studies as potential biomarkers for discrimination of bTB+ and bTB− cattle.

We therefore trained a total of eight ML models using 1,311 DEGs identified in the consolidated training set encompassing animals from all four bTB datasets (**Fig. 3A**). Using a five-fold cross validation framework for hyperparameter tuning, we observed high average AUROC values > 0.90 for the penalised logistic regression models (ENET, LASSO, and RIDGE), and the MLP, RF and RF-ET models, which generalised well and achieved comparable performance in the testing dataset (**Fig. 3C**, **Fig. 3D**), indicating that the PB transcriptome can accurately discriminate bTB+ and bTB− cattle. We also observed a trend whereby the best performing models yielded higher AUROC estimates in the experimentally infected compared to the naturally infected groups; however, this difference was not significant (*P* = 0.64, **Fig. 4E**). This observation may indicate that the inferred RNA biosignatures are being driven by an innate immune-biased signature, as the transcriptional profiles of animals recently experimentally infected with *M. bovis* exhibited perturbed innate immune response pathways; however, it may also be partially attributable to variations in the *M. bovis* infectious dose between the experimentally and naturally infected animals (**Fig. 2C**). This former explanation is supported by the observation that all experimentally infected animals from the MCL21-BTB dataset could be classified as bTB+ when optimum thresholds were selected to yield sensitivity values ≥ 0.90 in the testing set (**Fig. 3D**, **Fig. 3F**, **Table S10**). Moreover, the best-performing model (ENET), which achieved a sensitivity and specificity of 0.941 and 0.853, respectively, used expression information from 273 (20.8%) out of all input DEGs to determine the probability that an animal was bTB+. This biosignature was composed of key innate immune response genes, including *TLR4*, *MX1*, *IFNGR1*, *IFITM3*, *GBP2, GBP4*, *TNFAIP2*, *IFI6*, and *UBASH3A*; these genes have been shown to have pivotal roles in the host response to mycobacterial infections or in transcriptional biosignatures for hTB using whole blood (*50–55*).

We also generated three infection *z*-scores derived from three gene sets that were identified with a GFS algorithm (*28, 29, 42*), which resulted in a 13-gene set signature, a 17-gene set signature, and a combined 30-gene set signature (**Fig. 4A**, **Fig. 4B**). These three gene sets achieved high discrimination between bTB− and bTB+ cattle in the training set with AUROC values ≥ 0.96 across the five folds (**Fig. 4B**, **Fig. 4C**). For the training set, these gene sets achieved slightly lower but highly accurate discriminatory power between bTB+ and bTB− cattle, with all combinations achieving an AUROC > 0.85. The most discriminatory gene that contributed to the infection *z*-score identified in the 13-gene set was *SATB1*, which has previously been shown to be upregulated in macrophages upon infection with virulent *M. tuberculosis*, contributing to enhanced bacterial survival through repression of NADPH oxidase 2 (NOX2 – the catalytic core of NADPH oxidase encoded by *CYBB*) with a concomitant downregulation of bactericidal reactive oxygen species (ROS) production (*56*). The *PTK2* gene encoding the focal adhesion kinase (FAK) protein was identified as the most discriminatory gene in the 17-gene set (**Fig. 4A**). The concentration of FAK in macrophages infected with *M. tuberculosis* has been shown to decrease in a time-dependent manner orchestrated by the bacterium through inhibition of ROS production, culminating in increased macrophage necrosis and uncontrolled replication of *M. tuberculosis* (*57*). Other genes identified by the two GFS iterations included *NR4A2*, a transcription factor that regulates macrophage polarity from an M1 phenotype to an M2 phenotype (*58*), and which was previously identified as being upregulated in monocytes derived from non-infected household contacts heavily exposed to *M. tuberculosis* in comparison to monocytes isolated from individuals latently infected with *M. tuberculosis* (*59*); *CCR5*, which was upregulated in macrophages infected with *M. tuberculosis* resulting in enhanced IL-10 production (*60*); and *UBASH3A* described previously. These results indicate that ML models and the GFS approach have identified genes expressed in PB that can effectively discriminate between bTB+ and bTB− cattle and that play mechanistic roles in mycobacterial infection biology and host-pathogen interaction.

Analysis of ML models for other infectious diseases revealed variable discrimination between infected and control non-infected animals within each dataset. For example, in the ALO19-MAP dataset, the RIDGE classifier was the best performing model, achieving an AUROC of 0.667 (95% CI: 0.01–1.00) indicating poor to moderate discriminatory capacity (**Fig. 5A**). Although infection with MAP upregulated the expression of key immune-related genes such as *CXCL8* and *IFI27*, and perturbed common biological pathways observed in the bTB dataset, such as *Defence response* and *Immune response* (*39*), these results indicate that MAP induces a different PB transcriptional profile in cattle compared to *M. bovis*, which resulted in the observed suboptimal performance of ML models in this dataset.

Unlike MAP infection, we observed substantially higher discriminatory power between BoHV-1− and BoHV-1+ cattle, respectively, with the ENET model achieving high discriminatory capacity (AUROC = 0.931; 95% CI: 0.79−1.00) followed closely by the RF, RIDGE, LASSO, RF−ET and MLP models, which achieved AUROC values ranging from 0.847−0.917 (**Fig. 5B**). Additionally, we observed even higher discrimination between BRSV− and BRSV+ animals, ranging from perfect discrimination for the LASSO, MLP and RF-based models (AUROC = 1.00) to excellent performance for the RIDGE and ENET models with these two models each achieving AUROC estimates of 0.986 and 0.944, respectively. These findings suggest that infection with either BoHV-1 or BRSV induces transcriptional perturbations in PB of cattle that are similar but not identical to those observed after infection with *M. bovis*. These observations are supported by the biological pathways perturbed by DEGs in the BoHV-1 and BRSV datasets reported by (*41*) and (*40*), respectively. For the BoHV-1 dataset, SIPs included the *NOD*-*like receptor signalling* pathway, the *Defence response to virus* GO term, the *Interferon signalling* Ingenuity Pathway Analysis (IPA^®^) term, and the *Regulation of viral process* GO term (*41*). For the BRSV dataset, SIPs included the *Defence response to virus*, *Regulation of viral life cycle*, and *Innate immune response* GO terms (*40*). We also observed somewhat comparable variability across datasets in terms of discriminatory power between non-infected and infected cattle for the three gene sets inferred from the GFS algorithm (13-gene set, 17-gene set, and 30-gene set). The three gene sets achieved poor discrimination in the MAP dataset with AUROC values ranging between 0.545−0.667, extremely high discrimination in the BoHV-1 (AUROC range: 0.861–1.00) and extremely poor to excellent discriminatory power in the BRSV cohort (AUROC range: 0.639–0.972) (**Fig. 5E**). These results indicate that the gene sets inferred using the GFS algorithm closely resemble peripheral transcriptional perturbations of these genes induced by BoHV-1 but are distinct from those observed in MAP- and BRSV-infected cattle compared to healthy controls.

Lastly, we evaluated the ability of all ML models and GFS gene sets to discriminate between bTB+ animals and MAP+, BRSV+, and BoHV-1+ cattle as alternative disease-positive controls. We observed varying discriminatory power across all ML models, and gene sets with AUROC values ranging from 0.367 for the unpenalised logistic regression model to 0.868 for the MLP model (**Fig. 5F**). The second and third-ranked classifiers were the ENET classifier and the 30-gene set signature, which achieved AUROC estimates of 0.826 and 0.819, respectively. These results are noteworthy as the ENET classifier and the 30-gene set signature both achieved a specificity of 0.853 and sensitivity of ≥ 0.853 for the bTB testing group (**Fig. 3F** and **Fig. 4F**). The performances of these classifiers observed here indicate that they both have the capacity to differentiate between bTB+ and bTB− cattle effectively and are also robust at discriminating between bTB+ animals and cattle infected with other bacterial and viral pathogens.

In current bTB-endemic countries, control and eradication of the disease are primarily underpinned by test-and-slaughter programmes, in which infected animals are identified and removed from the herd (*7, 61*). The *in vivo* SICTT diagnostic is used as a population screening tool for bTB prevalence due to its high specificity (> 0.99 in Britain), and positive predictive value (PPV – the probability that a positive result is truly positive; > 0.91 in Britain) (*26*). In addition, the PPV improves as disease prevalence rises because confidence increases that a positive result is a true positive (*62, 63*). The ancillary *in vitro* IGRA test is typically employed when ≥ 5 SICTT reactor animals in a herd are identified and is interpreted in parallel with the SICTT to maximise the detection of infected cattle in these high-risk herds (*64–66*). In this regard, the sensitivity of the combined SICTT and IGRA tests is estimated at 93% (*20*).

The SICTT and IGRA tests are based on cell-mediated immunological responses and, as such, are limited to the detection of animals at least +2 wpi in the case of the IGRA test (*21*), and between +3 and +6 wpi for the SICTT (*67*). Although the sample size is limited in the testing set, our ENET model and 30-gene set signature correctly classified experimentally infected animals as being bTB+ at +1 and +2 wpi, respectively (**Fig. 3F**, **Fig. 4F**), indicating that these classifiers have different diagnostic properties compared to the SICTT and IGRA test; these observations suggest that a third diagnostic capable of detecting early disease cases, may increase the ability to identify all bTB+ animals in a herd and contribute significantly to control and eradication programmes. Notwithstanding these encouraging results, it is important to emphasise that to validate these observations, a much larger PB transcriptomics study using cattle experimentally infected with *M. bovis* at several early post-infection time points will be required.

While the bTB+ animals at +1 wpi and +2 wpi were correctly classified as such in the testing set by the ENET and 30-gene set classifiers, the two bTB+ animals that were consistently misclassified as bTB− based on the selected probability threshold values for these two models were T059 and T062, both of which were derived from the OGR25-BTB dataset (**Table S9** and **S16**). Interferon-γ release assay test results demonstrated that T059 yielded a positive result, whereas T062 produced a negative result for this diagnostic (*31*). These results indicate that while the ENET and 30-gene set classifiers, respectively, augment current bTB diagnostics through correctly classifying animals infected with *M. bovis* ≤ +2wpi as bTB+, they illustrate that training alternative PB transcriptomic models using data from SICTT+/IGRA−/culture+ animals or SICTT−/IGRA−/culture+ animals, respectively, is necessary, and represents a parallel approach to strengthen current bTB testing strategies further and to address the persistent issue of residual infection in bTB breakdown herds (*68*).

Similar to the *in vitro* IGRA test, the ENET classifier and infection *z*-score thresholds from the 30-gene set signature can be readily adjusted in high-risk herds where test specificity is less of an issue, because maximisation of sensitivity in these populations can aid in removing all infected animals, which will substantially improve control and eradication efforts (*20*). However, a larger cohort of naturally infected animals will be required to validate the findings and diagnostic estimates obtained in this study. This may be achieved through additional high-throughput transcriptomic analysis, which has seen dramatic cost reductions over recent years. Also, in addition to conventional bulk RNA-seq, targeted transcriptomic methods could also be used (*69, 70*).

Our analyses have demonstrated that machine learning classifiers trained on PB/PBL transcriptomics data can robustly distinguish bTB+ from bTB− cattle, and bTB+ cattle from cattle infected with other bacterial and viral pathogens. The outputs from this work highlight the potential of RNA-based biosignatures to complement and augment existing diagnostics in countries where bTB disease is endemic. Further validation and development of these classifiers into rapid diagnostic tools could enhance bTB control and eradication programmes in endemic regions.

## Materials and Methods

### Data acquisition and preprocessing

For this study, publicly available raw RNA-seq data were obtained from six experiments investigating transcriptional differences in peripheral blood (PB) or peripheral blood leukocyte (PBL) samples between control non-infected cattle and animals infected naturally or experimentally with: 1) *M. bovis* (bTB+), encompassing three datasets reported by (*35*) (MCL14-BTB), (*38*) (WIA20-BTB), and (*36*) (MCL21-BTB); 2) Bovine respiratory syncytial virus (BRSV) generated by (*40*) (JOH21-BRD); 3) Bovine herpes virus (BoHV-1) analysed by (*41*) (ODO23-BRD) and; 4) *M. avium* ssp. *paratuberculosis* (MAP) reported by (*39*) (ALO19-MAP) (**Table 1**). These raw datasets were downloaded using fastq-dl (v. 2.0.4). Lastly, an additional pre-processed RNA-seq count matrix file, derived from *n* = 63 control (bTB−) and *n* = 60 (bTB+) cattle from (*31*) (OGR25-BTB), was downloaded from the Gene Expression Omnibus (GEO) database using the BioProject accession number <u>GSE255724</u> (**Table 1**).

The quality of the raw RNA-seq data from MCL14-BTB, WIA20-BTB, MCL21-BTB, JOH21-BRD, ODO23-BRD and ALO19-MAP datasets was assessed using fastqc (v. 0.11.9) and multiqc (v. 1.25.1) (*71*). Illumina adaptors were trimmed using trimmomatic (v. 0.39) (*72*) and raw reads were further filtered to retain reads ≥ 36 bp and to remove trailing bases with a phred-scale quality score < 30. The trimmed RNA-seq reads were aligned to the ARS-UCD1.2 bovine reference genome (*73*) using STAR (v. 2.7.1) (*74*). The reference genome index was generated using the *genomeGenerate* command with the *Bos taurus* ARS-UCD1.2 Ensembl annotation file (v. 110) specified in addition to the--*sjdbOverhang* 99 parameter. Read counts for each gene were quantified using featureCounts (v. 2.0.0) (*75*).

### Variant calling from RNA-seq data and population genomics analyses

To call variants directly from the RNA-seq data, we followed the Pig Genotype Tissue Expression (GTEx) Consortium’s workflow (https://github.com/FarmGTEx/PigGTEx-Pipeline-v0/blob/master/02_RNA-Seq/03_SNP_calling.smk) (*76*) and used the Genome Analysis Tool Kit (GATK) (v. 4.3.0.0) (*77*). Note: imputed whole-genome sequence data (WGS) comprising a total of 3,866,506 filtered autosomal SNPs (minor allele frequency (MAF) ≥ 5%, non-significant deviation from Hardy-Weinberg Equilibrium (HWE) *P* ≥ 1 × 10^−6^, imputed dosage *R*^2^ ≥ 0.6) was available for the *n* = 123 animals used in the experiment conducted by (*31*) (https://zenodo.org/records/13752056).

Briefly, for the RNA-seq data derived from the time series experiments (*36, 38*), we first merged animal-specific BAM files across time points to increase read coverage for the variant call. Next, for all independent BAM samples across the RNA-seq-only experiments (*n* = 39), we applied the GATK Best Practices guidelines for variant discovery (*78, 79*). We subsequently filtered out low-quality variants using the following criteria: FisherStrand (FS) test > 30.0, QualByDepth (QD) < 2.0, read depth (DP) < 4.0; we also restricted our analysis to biallelic autosomal SNPs. We then merged the sample and study-specific VCF files using the BCFtools (v. 1.15.1) (*80*) *merge* command. Overall, this analysis yielded 163,589, 989,438, and 1,482,214 raw variants called in the MCL14-BTB, WIA20-BTB, and MCL21-BTB datasets, respectively.

We next intersected the above RNA-only VCF files with the filtered WGS data from (*31*) using the BCFtools *isec* command with options ‘*-*n +4’ and ‘-c none’ to retain a total of 31,312 SNPs called/present across all datasets. Next, we merged all datasets and filtered the 31,312 SNPs for those with a MAF > 5%, that did not deviate from HWE (*P* < 1 × 10^−6^), and that were not missing in more than 20% of samples. This procedure resulted 29,740 SNPs being retained. From these, we extracted all the samples from the MCL14BTB, WIA20-BTB, MCL21-BTB and OGR25-BTB datasets, respectively, into study-specific VCF files.

The filtered VCF files resulting from the workflow detailed above were loaded into R (v. 4.3.2) (*81*) with the SNPRelate (v. 1.34.1) toolset (*82*) using the *snpgdsVCF2GDS* and *snpgdsOpen* functions. A principal component analysis (PCA) was conducted on each SNP dataset using the *snpgdsPCA* function, and the eigenvectors corresponding to principal component 1 (PC1) and PC2 were plotted using ggplot2 (v. 3.4.4) (*83*). As a validation step, we also compared the eigenvector coordinates of PC1 and PC2 derived from the 29,740 SNPs in the OGR25-BTB dataset to the eigenvector coordinates for PC1 and PC2 from the same set of animals derived from 34,272 pruned genome-wide array SNPs initially reported by (*31*) using the Spearman correlation (*ρ*_spearman_).

### Intra-bTB dataset differential expression analysis

A DEA was conducted separately between the reactor (bTB+) and control (bTB−) animal groups for each bTB dataset using DESeq2 (v. 1.40.2) (*84*). For the generalised linear model, a design matrix was specified, which included the following covariates: age in months (where applicable), RNA-seq sequencing batch (where appropriate), and population genetic structure, in the form of PC1 and PC2 from the PCA of the RNA-seq SNPs for each dataset, with infection status selected as the variable of interest. The PC1 and PC2 covariates were included because the crossbred/multibreed nature of the animals in each study population should be incorporated in the DEA model design. Genes with raw expression counts ≥ 6 in at least 20% of samples were retained prior to the DEA. For the DEA, the null hypothesis was that the LFC between the bTB+ and bTB− groups, for the expression of a particular gene is exactly 0. To account for potential heteroscedasticity of LFCs, we implemented the approximate posterior estimation for generalised linear model coefficients (APEGLM) method (*85*) using the *lfcShrink* function. Genes with a Benjamini-Hochberg (BH) false discovery rate (FDR) adjusted *P*-value (*86*) (FDR-*P*_adj._) < 0.05 and a |LFC| > 0 were considered significantly DEGs.

For the OGR25-BTB and MCL2014-BTB datasets, a straightforward comparison between bTB− and bTB+ cattle was conducted. For the MCL21-BTB dataset, five comparisons were performed to compare the bTB+ animals at each time point (+1 wpi, +2 wpi, +6 wpi, +10 wpi, and +12 wpi) with the bTB− animals pre-infection (−1 wpi). For the WIA20-BTB dataset, the bTB+ animals at each time point (+4 wpi, +10 wpi) were compared to the combined *n* = 13 bTB+ and bTB− animals at 0 wpi; the “bTB+” animals at 0 wpi were sampled before experimental infection with *M. bovis* and are therefore designated as bTB−. The DE results were plotted as volcano plots using ggplot2 (v. 3.4.4) (*83*) and the intersections of DEGs among studies were visualised using the UpSetR R package (v. 1.4.0) (*87*).

### Functional enrichment analysis of differentially expressed genes

A functional enrichment analysis was conducted for DEGs inferred from each bTB dataset using the g:GOSt tool in g:Profiler (v. 0.2.2) (*88*). The organism selected was *Bos taurus*, and genes exhibiting increased, and separately decreased expression, were used to generate ordered input query lists based on the DEA FDR-*P*_adj._ values. Note: For the analysis of time series data, we aggregated results across time points, filtered for DEGs with a consistent direction of effect (either LFC > 0 or LFC < 0) across time points, and ordered the retained genes by DEA FDR-*P*_adj._ values. In addition, we followed best practice recommendations to account for tissue-specific sampling biases in gene set overrepresentation and functional enrichment analyses (*89*). Consequently, for the analysis of DEGs, the background set consisted of all expressed genes tested for differential expression across the bTB datasets. For analyses of our query DEG sets, we selected the gene ontology biological process (GO:BP) and cellular component (GO:CC) categories from the GO knowledgebase (*90*) in addition to the Kyoto Encyclopaedia of Genes and Genomes (KEGG) (*91*) and Reactome (*92*) resources. To identify significantly enriched/overrepresented pathways, a BH-FDR multiple testing correction was applied (FDR-*P*_adj._ < 0.05).

### Generation of training and testing sets

We randomly divided the complete bTB dataset (*n* = 230) into a training set (70%; *n* = 162: *n* = 69 (bTB−) and *n* = 93 (bTB+)) and a testing set (30%; *n* = 68: *n* = 34 (bTB−) and *n* = 34 (bTB+)), ensuring that the repeated measures of RNA-seq data derived from animals involved in the time series experiments were included in only one of the two sets to avoid overlap. For the training of ML models and evaluation of hyperparameters in the training set, we further divided the training set into five folds: Fold 1 (*n* = 35: *n* = 14 (bTB−) and *n* = 21 (bTB+)); Fold 2 (*n* = 36: *n* = 17 (bTB−) and *n* = 19 (bTB+)); Fold 3 (*n* = 32: *n* = 16 (bTB−) and *n* = 16 (bTB+)); Fold 4 (*n* = 34: *n* = 10 (bTB−) and *n* = 24 (bTB+)); and Fold 5 (*n* = 25: *n* = 12 (bTB−) and *n* = 13 (bTB+)). This procedure was implemented in a manner similar to the broad train/test splitting procedure, ensuring that repeated measures of RNA-seq data from animals in the time-series experiments were not present across folds.

### Differential expression analysis on bTB training data

Within the entire training dataset, a DEA was conducted between the reactor (bTB+) and control (bTB−) animal groups using DESeq2 (v. 1.40.2) accounting for age in months binned into four categories (1–6 months, 6–12 months, 12–24 months, and > 24 months); study/batch (five levels: two for the OGR25-BTB dataset and one each for the MCL14-BTB, WIA20-BTB, and the MCL21-BTB datasets, respectively); and PC1 and PC2 derived from the PCA of the pruned RNA-seq called SNPs across all samples in the training set as covariates, with condition (bTB+ or bTB−) set as the variable of interest.

For this DEA, the null hypothesis was that the LFC between the bTB+ and bTB− groups, for the expression of a particular gene, is exactly 0. Similar to the intra-dataset DE analysis, to account for potential heteroscedasticity of LFCs, we implemented the approximate posterior estimation for generalised linear model coefficients (APEGLM) method using the *lfcShrink* function. Genes with an FDR-*P*_adj._ < 0.05 and a |LFC| > 0 were considered to be significant DEGs. For the identification of features for inclusion in the development of ML classifiers, we *post-hoc* filtered DEGs and retained those with a mean median-of-ratios normalised expression value (*84*) > 100 (*29*).

### Training of integrated classifiers on bTB training data

Machine learning (ML) model training and hyperparameter grid search were performed using R (v. 4.3.2) (*81*). All ML classification models were generated using the variance-stabilised transformed (VST) counts for the *post-hoc* filtered DEGs obtained using the *varianceStabilizingTransformation* function from the DESeq2 R package as input in the caret R package (v. 6.0-90) (*93*) with results evaluated using the pROC package (v. 1.18.4) (*94*) with performance metrics including sensitivity, specificity, positive likelihood ratio (LR+), negative likelihood ratio (LR−) and the area under the receiver operating characteristic curve (AUROC) (*95*), extracted, tabulated and visualised using ggplot2 (v. 3.4.4) where appropriate. The classification models and hyperparameters evaluated in the training set were: unpenalised generalised logistic model (GLM; no hyperparameters); generalised logistic regression with ridge feature selection (RIDGE; hyperparameters: α = 0, λ = [10^-5^–10^0^, for 10,000 values]); generalised logistic regression with least absolute shrinkage and selection operator feature selection (LASSO; hyperparameters: α = 1, λ = [range = 10^-5^–10^0^, for 10,000 values]); generalised logistic regression with elastic net feature selection (ENET; hyperparameters: α = [range = 0.05−0.95, in steps of 0.05], λ = [range = 10^-5^–10^0^, for 10,000 values]); random forest (RF; hyperparameters: mtry = [range = 2−35, in steps of 5], min.node.size = [range = 2−10, in steps of 1], num.trees = [501, 751, 1001, 1501, 3001, 5001], splitrule = “gini”]); random forest with the ExtraTrees algorithm (RF-ET; hyperparameters: mtry = [range = 5−100, in steps of 5], min.node.size = [range = 2−10, in steps of 1], num.trees = [501, 751, 1001, 1501, 3001, 5001], splitrule = “extratress”]); naïve Bayes (NB; hyperparameters: laplace = [in steps of 0 – 3 in steps of 0.1], adjust = [10^-1^−10^1^, 0.1], usekernel = TRUE); and a multi-layered perceptron (MLP; hyperparameters: size = [2−10, 1], decay = [range = 10^-9^−10^1^, in steps of 10*^n^* where *n* = [-9−1]). The confidence intervals of AUROC estimates were computed using the *ci* function from the pROC R package, which performed stratified bootstrap resampling 2000 times (*94, 96*).

### Greedy forward search strategy and generation of a bTB infection *z*-score

Previous studies have shown, across a wide range of diseases, including TB (*28, 29*), sepsis (*97*), and dengue fever (*43*), that a greedy forward search (GFS) algorithm can be used to generate a disease-specific “infection *z*-score” (*42*), which is derived from a panel of DEGs with high discriminatory power between cases and controls. The infection *z*-score is calculated as the mean of the VST normalised expression levels for positive DEGs (LFC > 0) minus the mean of the VST normalised expression levels for all negative DEGs (LFC < 0) (*29, 42*). In this study, the positive and negative DEGs, respectively, are derived from the comparison of bTB+ and bTB− cattle in the training set.

The GFS algorithm starts with a single gene exhibiting the best discriminatory power, represented as the average AUC (area under the curve) across the five folds used in the training set, and then at each subsequent step adds the gene with the best possible increase in the average AUC to the set of genes, until no further additions can increase the average AUC. Given substantial transcriptional heterogeneity across bTB datasets, we sought to generate a robust diagnostic signature set rather than aiming for extreme parsimony. Therefore, we ran the forward search a second time, such that once the first gene set had been identified, those genes were removed from the remaining pool and the forward search was run again. We subsequently generated a third combined list of 30 genes derived from the sets of genes identified in the first and second iterations, respectively.

### Assessment of classifier performance on bTB testing data

To normalise the testing data, we first froze the VST estimates of the training data using the *dispersionFunction* function from DESeq2 and subsequently applied these frozen VST estimates to the testing data using the same function. We then VST-normalised the testing data using the updated dispersion estimates with the *varianceStabilizingTransformation* function from DESeq2, setting the *blind* parameter to FALSE. Within the testing set, we extracted the VST counts of DEGs from the training set, then selected the hyperparameter-tuned models and gene sets from the GFS strategy for evaluation.

### Evaluation of classifier performance on other infectious diseases

We evaluated the performance of the hyperparameter-tuned models and inferred GFS gene sets using publicly available RNA-seq data for three other diseases, including Johne’s disease caused by *M. avium* spp. *paratuberculosis* (MAP) (*n* = 3 MAP−, *n* = 11 MAP+) (*39*) and bovine respiratory disease (BRD) caused by infection with either BoHV-1 (*n* = 6 BoHV-1−, *n* = 12 BoHV-1+) (*41*), or BRSV (*n* = 6 BRSV−, *n* = 12 BRSV+) (*40*), respectively.

We first evaluated the performance of each classifier in an intra-dataset manner, comparing its discriminatory power between control non-infected animals and cattle infected with each of the respective agents, to highlight common and divergent host peripheral transcriptional changes associated with *M. bovis*, MAP, BoHV-1, or BRSV infections. For each dataset, we followed the same procedure used for normalising the bTB testing dataset, which included applying the frozen VST estimates from the bTB training dataset to each external dataset via the *dispersionFunction* function from DESeq2, followed by obtaining the VST normalised counts using the *varianceStabilizingTransformation* function from DESeq2, setting the *blind* parameter to FALSE. We then extracted the VST counts for DEGs from the bTB training set and selected the hyperparameter-tuned models and gene sets from the bTB training set GFS strategy for evaluation.

Finally, we evaluated the specificity of each classifier by assessing its discriminatory power between bTB+ animals (*n* = 34) in the testing set and MAP+ (*n* = 11), BRSV+ (*n* = 12), and BoHV-1+ (*n* = 12) animals. To do this, we extracted and integrated all bTB+ animals from the testing set and all MAP+, BRSV+, and BoHV-1+ animals from the three non-bTB datasets and set the *TB_status* variable for all bTB+ animals to *Infected* and for all other animals (MAP+, BRSV+, and BoHV-1+) to *Control.* We then normalised the expression matrix using the same procedure as for the testing set and the non-bTB dataset evaluations.

## Acknowledgements

We thank Michael McDonald for assistance with animal handling and blood collection.

## Funding

Research Ireland Centre for Research Training in Genomics Data Science grant no. 18/CRT/6214 (JFO).

Science Foundation Ireland Investigator Programme Award SFI/08/IN.1/B2038 (DEM and SVG).

Science Foundation Ireland Investigator Programme Award SFI/15/IA/3154 (DEM and SVG).

University College Dublin–University of Edinburgh Strategic Partnership in One Health (DEM, SVG, JGDP, and ELC).

Department of Agriculture, Food and the Marine (DAFM) project grant no. 17/RD/US-ROI/52 (DEM).

Department of Agriculture, Food and the Marine (DAFM) project grant no. 2023RP902 (DEM).

European Network on Livestock Phenomics (EU-LI-PHE) COST Action grant no. CA22112 (DEM, JFO, AK, ELC, and HP)

Fulbright-Teagasc Ireland-USA grant (JFO).

## Author contributions

Conceptualization: JFO, KGM, EG, ICG, SVG, CSG, DEM

Funding Acquisition: DEM, CSG, ICG, SVG, JGDP, ELC, HP, AK, JFO.

Methodology: JFO, AI, CNC, JGDP, SLFO, VR, HP, JAB, GPM, AK, TJH, EG, CSG, DEM.

Investigation: JFO, AI, EG, SVG, CSG, DEM.

Supervision: ICG, SVG, CSG, DEM.

Writing—original draft: JFO DEM and CSG.

Writing—review and editing: JFO, AI, GPM, CNC, JP, HP, KGM, SVG, CSG, DEM.

## Competing interests

Authors declare that they have no competing interests.

## Data and materials availability

All data needed to evaluate the conclusions in the paper are present in the paper and/or the Supplementary Materials. These raw and pre-processed RNA-seq datasets used in this study are publicly available under the following accession numbers: 1) OGRA25-BTB (<u>GSE255724</u>) (*31*); 2) MCL14-BTB (<u>PRJNA257841</u>) (*35*); 3) MCL21-BTB (<u>PRJEB27764</u> (bTB−) an <u>PRJEB44470</u> (bTB+)) (*36*); 4) WIA20-BTB (<u>PRJNA600004</u>) (*38*); 5) ALO19-MAP (<u>PRJNA565369</u>) (*39*); 6) JOH21-BRD (<u>GSE152959</u>) (*40*); and 7) ODO23-BRD (<u>GSE199108</u>) (*41*). Imputed and filtered WGS data from (*31*) are available at Zenodo (https://zenodo.org/records/13752056). Raw genotype information derived from the RNA-seq variant call for bTB studies described here is also available at Zenodo (https://doi.org/10.5281/zenodo.16887104).

The computer code and scripts used in this study, in addition to the trained models are available at the following GitHub link: https://github.com/jfogrady1/ML4Tb.

## Notes

https://www.ncbi.nlm.nih.gov/geo/query/acc.cgi?acc=GSE255724

https://zenodo.org/records/13752056

https://doi.org/10.5281/zenodo.16887104

### Competing Interest Statement

The authors have declared no competing interest.

### Summary of Updates

The title has been modified, and the abstract has been shortened. In addition, minor changes have been made to the manuscript text.

