## Supplementary Figures for "A new diagnostic modality for bovine tuberculosis: accurate and robust classification of infected cattle using transcriptomics and machine learning"

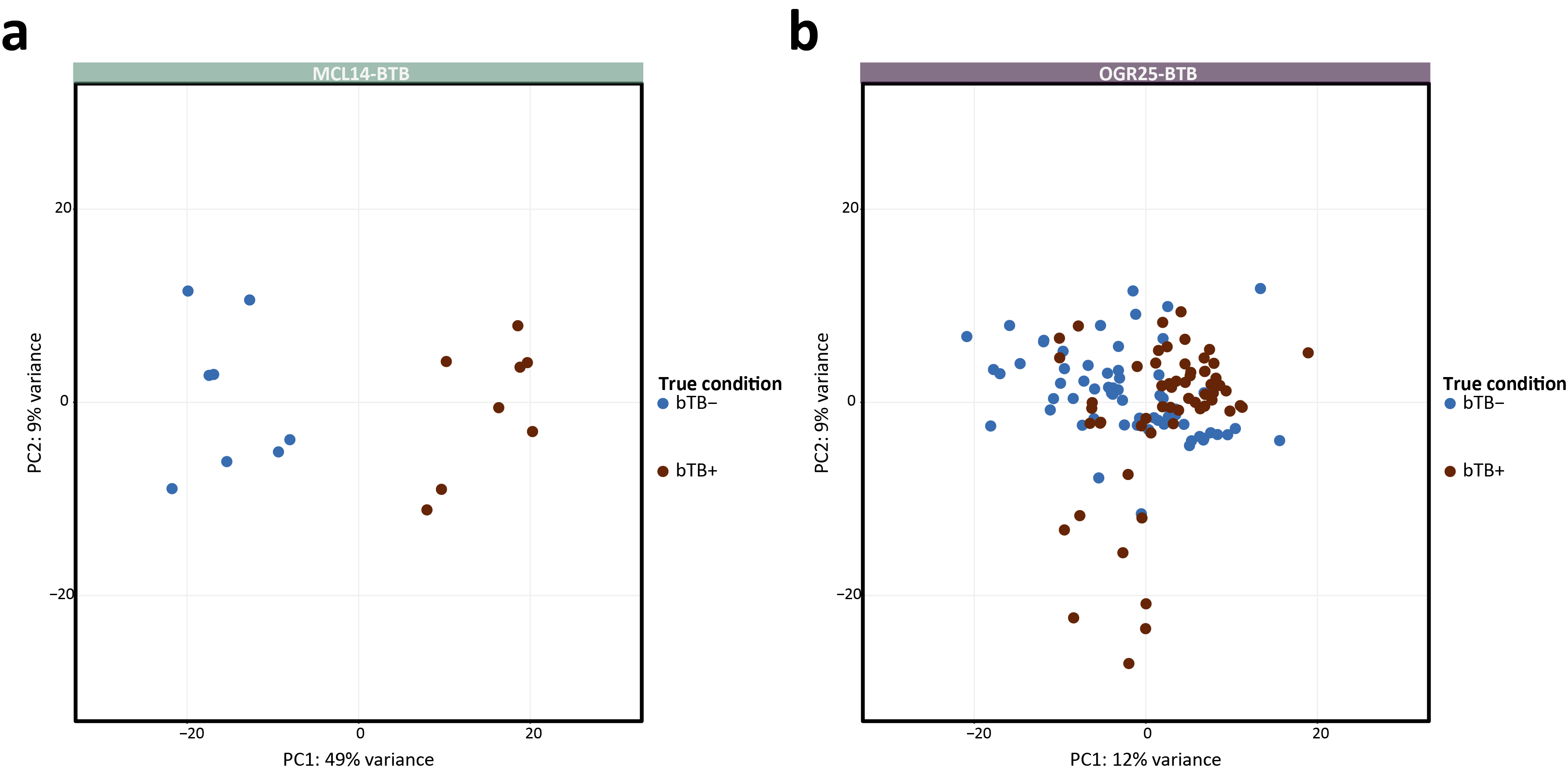

**Fig. S1**: Principal component analysis (PCA) of the top 750 most variable genes identified in (**a**) all *n* = 16 RNA-seq samples from the MCL14-BTB dataset and (**b**) all *n* = 123 RNA-seq samples from the OGR25-BTB dataset after variance stabilising transformation (VST) using DESeq2 (Love *et al.* 2014). Principal components 1 and 2 (PC1 and PC2) are plotted. Animal data points are coloured based on their experimental condition.

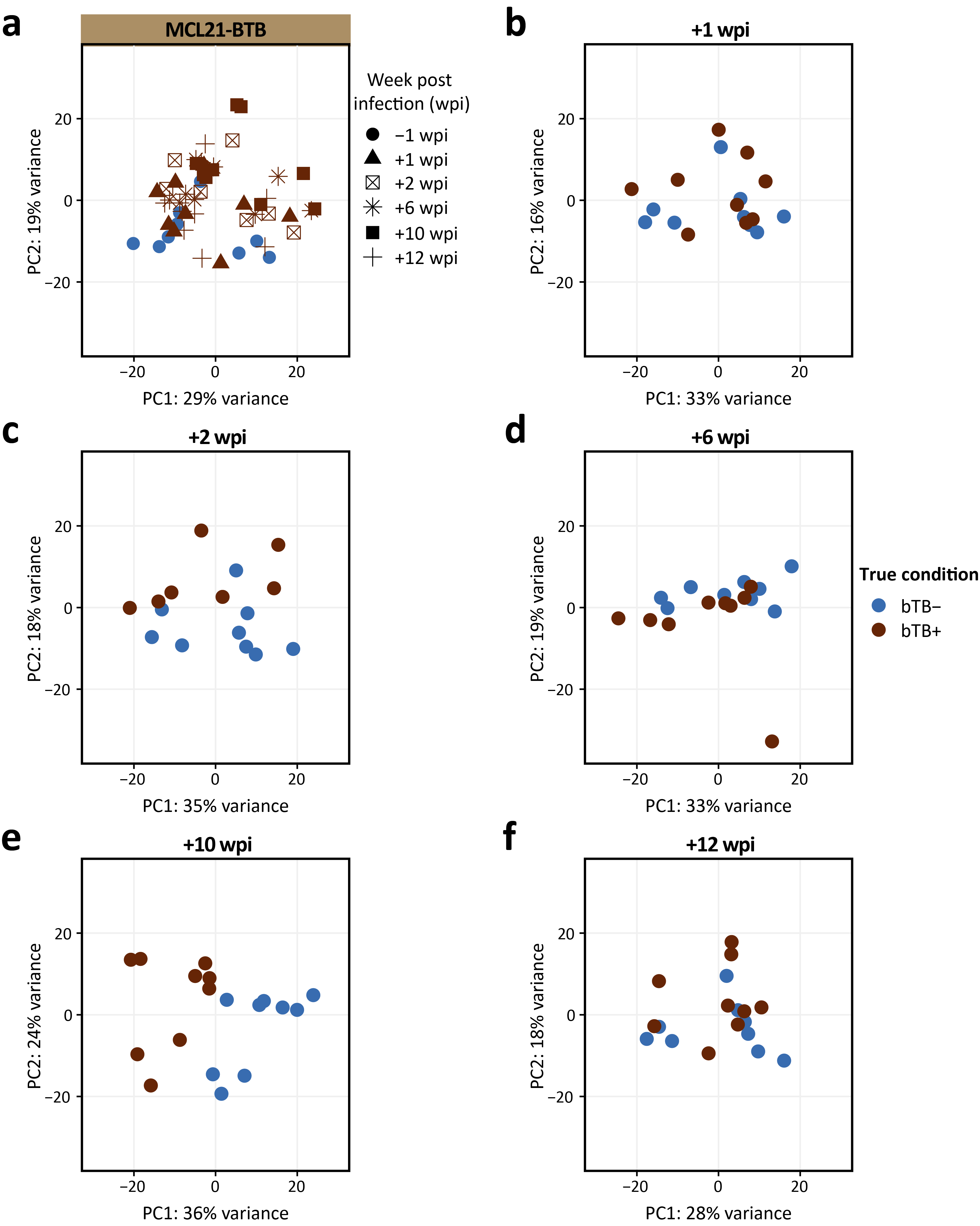

**Fig. S2**: Principal component analysis (PCA) of the top 750 most variable genes identified for (**a**) all *n* = 52 RNA-seq samples from the MCL21-BTB dataset after variance stabilising transformation (VST) using DESeq2. Principal components 1 and 2 (PC1 and PC2) are plotted. Animal data points are coloured by experimental condition and shaped according to their sampling time point, measured in weeks post-infection (wpi). (**b**) PCA of animals sampled at +1 wpi compared to animals sampled at −1 wpi. (**c**) PCA of animals sampled at +2 wpi compared to animals sampled at −1 wpi. (**d**) PCA of animals sampled at +6 wpi compared to animals sampled at −1 wpi. (**e**) PCA of animals sampled at +10 wpi compared to animals sampled at −1 wpi. (**f**) PCA of animals sampled at +12 wpi compared to animals sampled at −1 wpi.

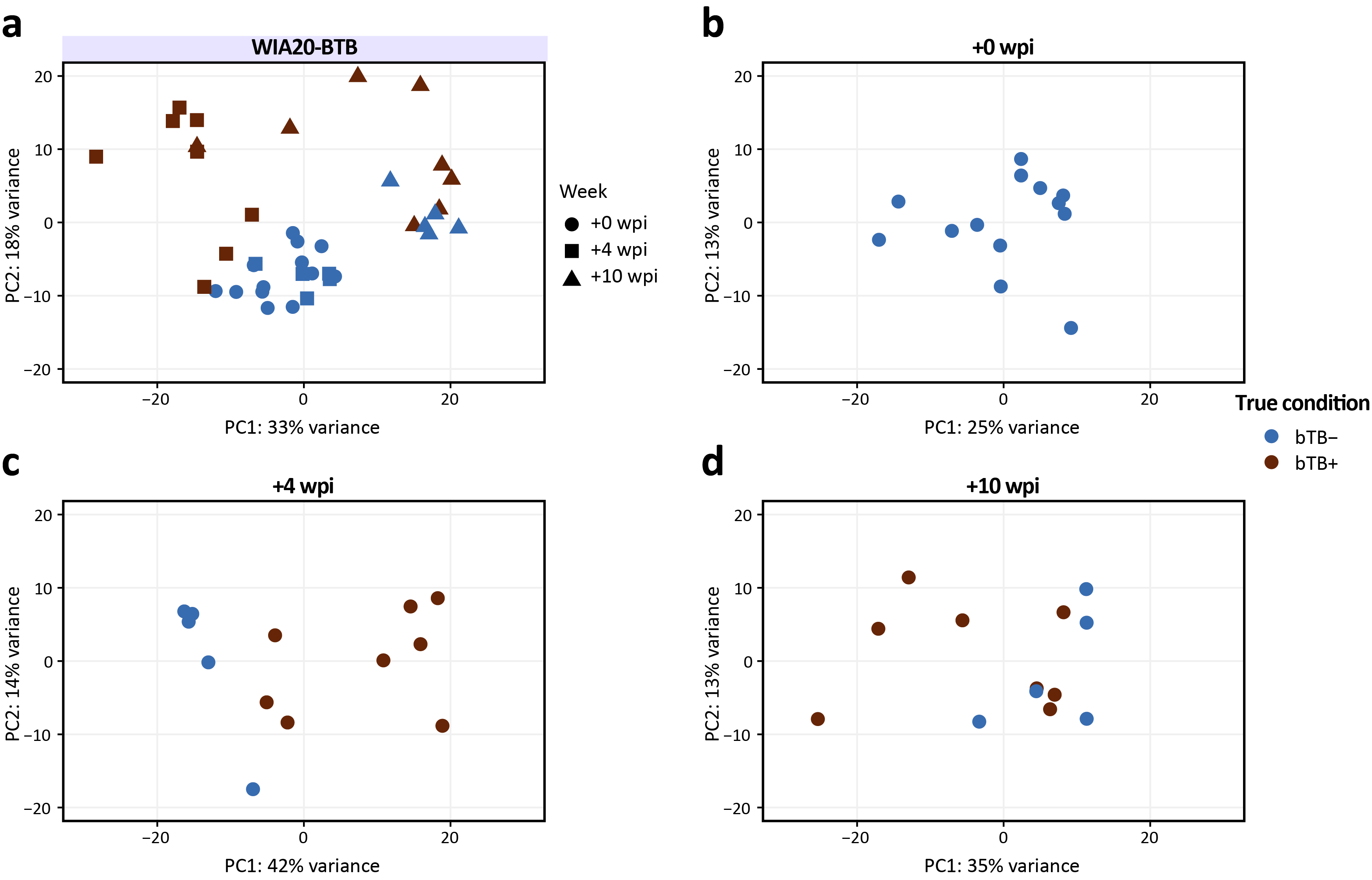

**Fig. S3**: Principal component analysis (PCA) of the top 750 most variable genes identified for (**a**) all *n* = 39 RNA-seq samples from the WIA20-BTB dataset after variance stabilising transformation (VST) using DESeq2. Principal components 1 and 2 (PC1 and PC2) are plotted. Animal data points are coloured by experimental condition and shaped according to their sampling time point, measured in weeks post-infection (wpi). (**b**) PCA of animals sampled at +0 wpi. Note: all *n* = 8 animals experimentally infected with *M. bovis* were sampled prior to inoculation and are therefore considered control (bTB−) animals at this time point. (**c**) PCA of animals sampled at +4 wpi compared to animals sampled at 0 wpi. (**d**) PCA of animals sampled at +10 wpi compared to animals sampled at 0 wpi.

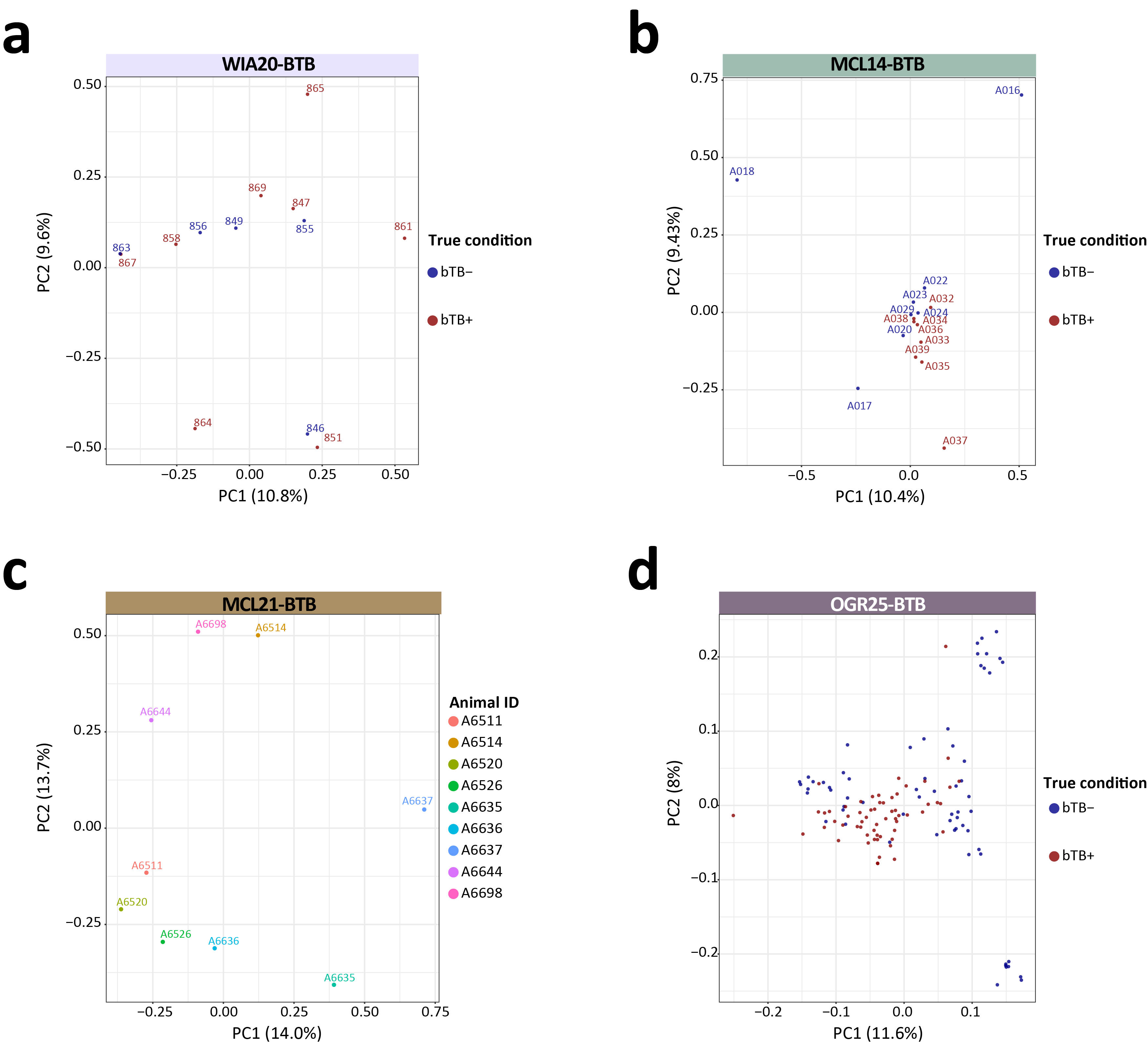

**Fig. S4**: (**a**) Genotype principal component analysis (PCA) of 29,740 genome-wide SNPs called from the RNA-seq data in the WIA20-BTB dataset. Animals are coloured based on whether they were experimentally infected with *M. bovis* or not. (**b**) Genotype PCA of 29,740 genome-wide SNPs called from the RNA-seq data in the MCL14-BTB dataset. Animals are coloured based on their experimental designation. (**c**) Genotype PCA of 29,740 genome-wide SNPs called from the RNA-seq data in the MCL21-BTB dataset. Animals are coloured according to their assigned ID. (**d**) Genotype PCA of 29,740 genome-wide SNPs called from the OGR25-BTB dataset. Animals are coloured based on their experimental designation. For all panels, principal components 1 and 2 (PC1 and PC2) are plotted.

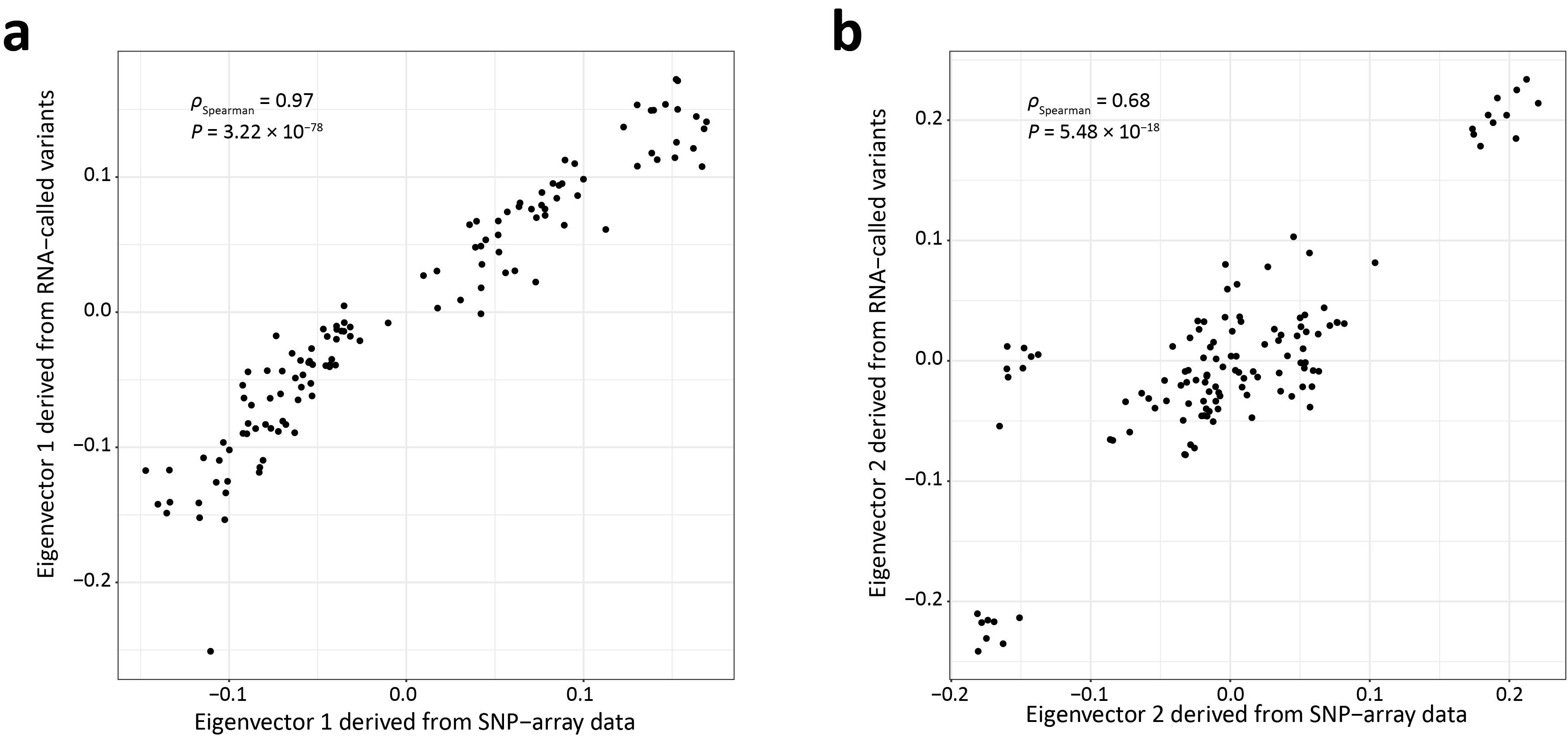

**Fig. S5**: (**a**) Spearman correlation (*ρ*) between eigenvector 1 coordinates for all *n* = 123 animals from the OGR25-BTB dataset derived from a principal component analysis (PCA) of 29,740 genome-wide SNPs and eigenvector 1 derived from a PCA of 34,272 pruned genome-wide SNP array data reported for the same set of animals in O'Grady *et al.* (2025) . (**b**) Same as **a,** but for eigenvector 2 coordinates.

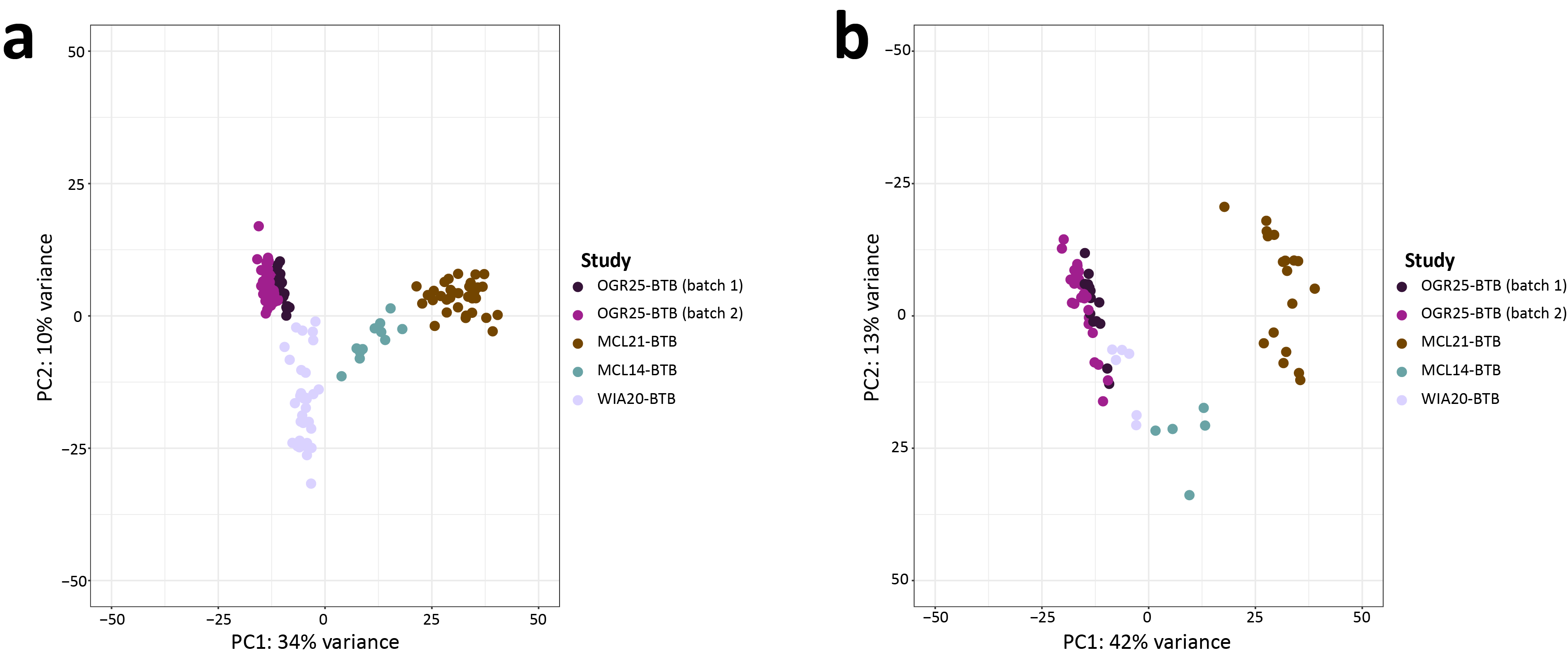

**Fig. S6**: (**a**) Principal component analysis (PCA) of the top 750 most variable genes identified in the training set after variance stabilising transformation (VST) using DESeq2. Principal components 1 and 2 (PC1 and PC2) are plotted. Animals are coloured based on their designated study. (**b**) A PCA of the top 750 most variable genes identified in the testing set after VST using DESeq2. PC1 and PC2 are plotted, and animals are coloured based on their designated study.

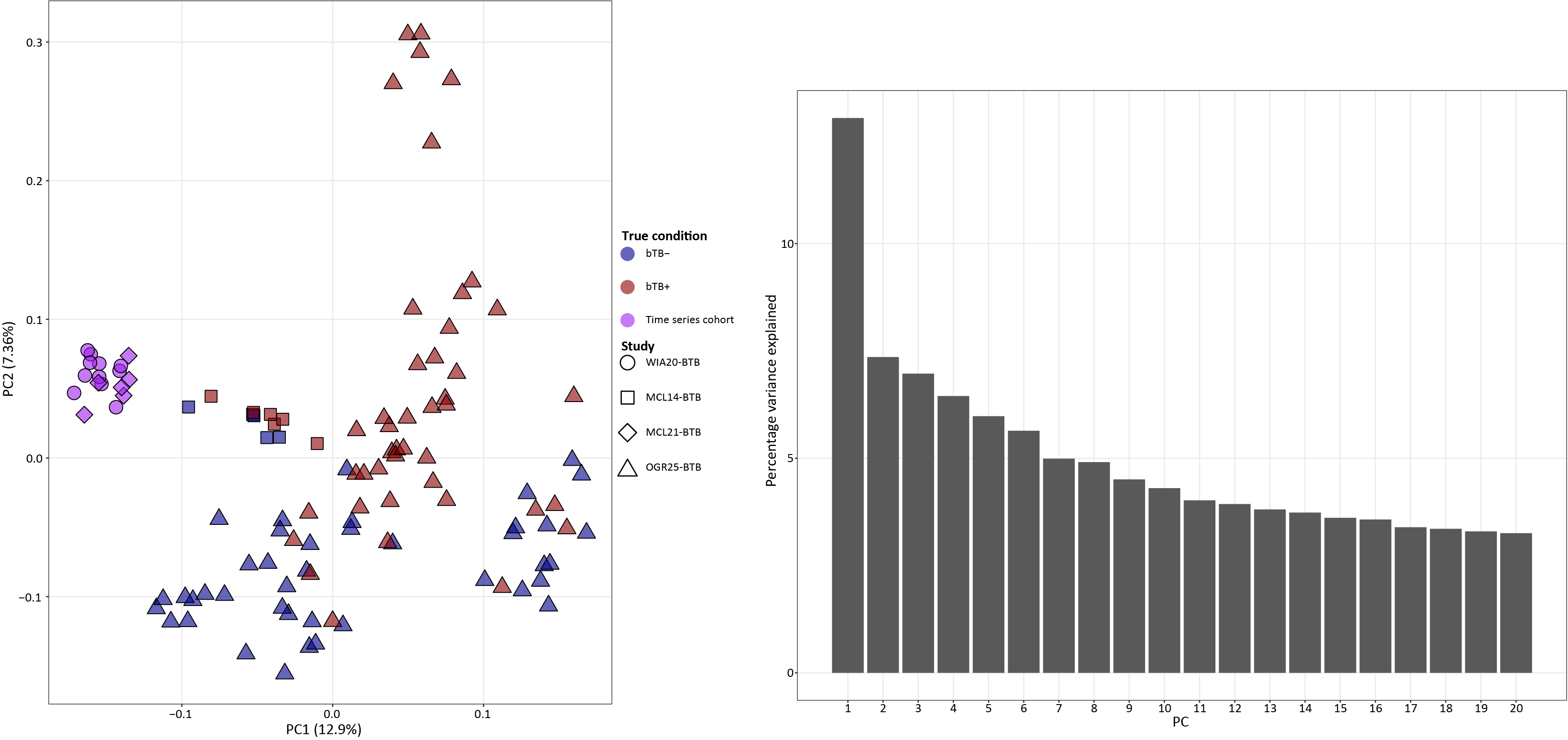

**Fig. S7**: Genotype principal component analysis (PCA) of 29,740 genome-wide SNPs identified in the training set. Principal components 1 and 2 (PC1 and PC2) are plotted. Animals are coloured depending on their experimental designation or if they are derived from a time series experiment. Animals are shaped based on their designated study. The barplot shows the proportion of variance explained by each of the top 20 PCs.

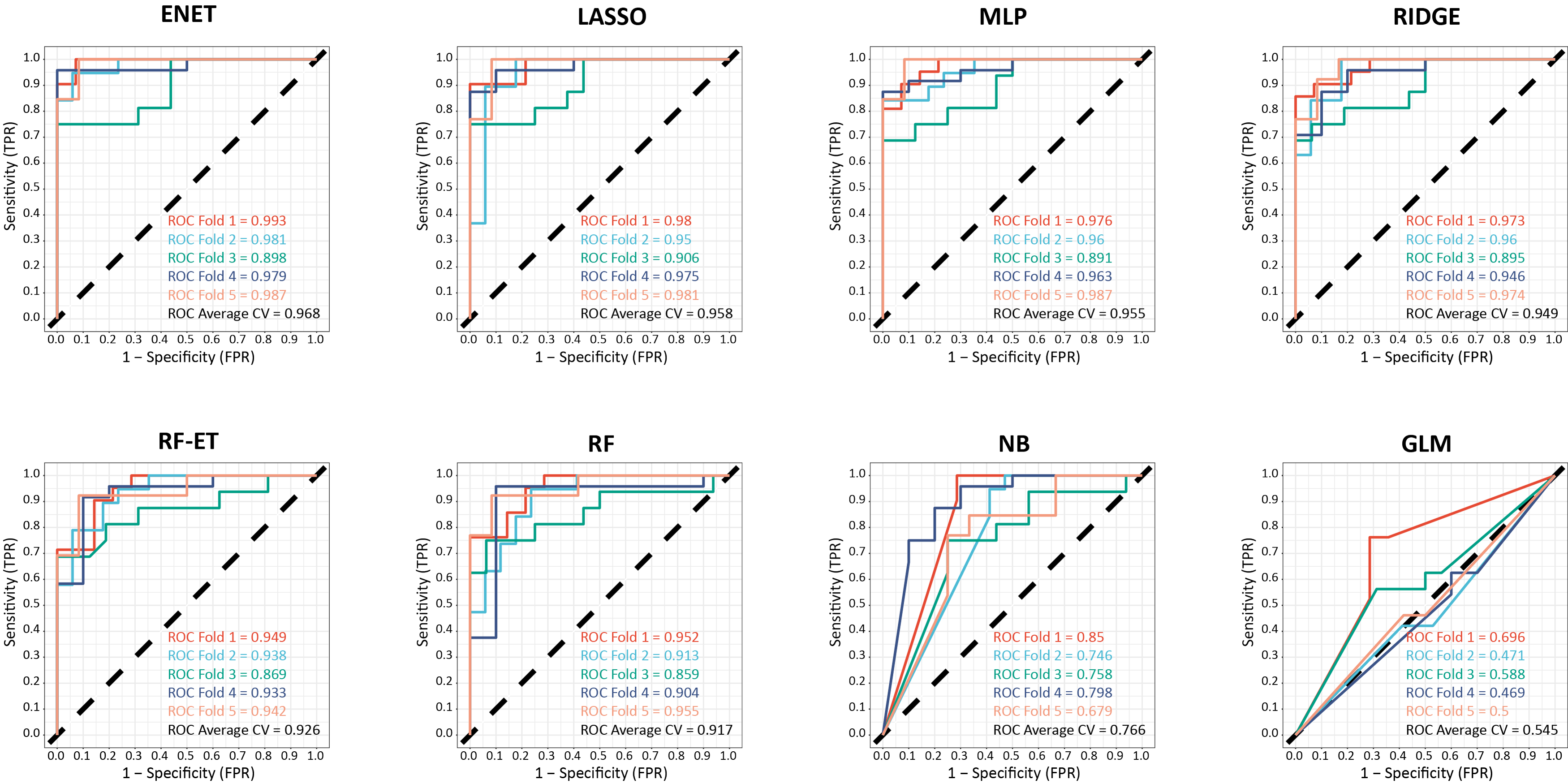

**Fig. S8**: Area under the receiver operating characteristic curve (AUROC) plots for eight machine learning (ML) models in each cross-validation fold in the training set. ENET, elastic-net; MLP, multi-layered perceptron; LASSO, least absolute shrinkage and selection operator; RIDGE, ridge-penalised regression; RF-ET, random forest (extra trees); RF, random forest; NB, naïve Bayes; and GLM, generalised unpenalized logistic regression model.
